# Structures and functions linked to genome-wide adaptation of human influenza A viruses

**DOI:** 10.1101/367169

**Authors:** Thorsten R. Klingen, Jens Loers, Stephanie Stanelle-Bertram, Gülsah Gabriel, Alice C. McHardy

**Affiliations:** Department for Computational Biology of Infection Research, Helmholtz Center for Infection Research, Braunschweig, Germany; German Center for Infection Research (DZIF), Braunschweig, Germany; Heinrich Pette Institute, Leibniz Institute for Experimental Virology, Hamburg, Germany; University of Veterinary Medicine, Hannover, Germany

**Keywords:** influenza A virus, positive selection, graph-cut, adaptation

## Abstract

Human influenza A viruses elicit short-term respiratory infections with considerable mortality and morbidity. While H3N2 viruses circulate since almost 50 years, the recent introduction of pH1N1 viruses presents an excellent opportunity for a comparative analysis of the genome-wide evolutionary forces acting on both subtypes. Here, we inferred patches of sites relevant for adaptation, i.e. being under positive selection, on eleven viral protein structures, from all available data since 1968 and correlated these with known functional properties. Overall, pH1N1 had more patches than H3N2 viruses, especially in the viral polymerase complex, while antigenic evolution is more apparent for H3N2 viruses. In both subtypes, NS1 had the highest patch and patch site frequency, indicating that NS1-mediated viral attenuation of host inflammatory responses is a continuously intensifying process, elevated even in the longtime-circulating subtype H3N2. We confirmed the resistance-causing effects of two pH1N1 changes against oseltamivir in NA activity assays, demonstrating the value of the resource for discovering functionally relevant changes. Our results represent an atlas of protein regions and sites with links to host adaptation, antiviral drug resistances and immune evasion for both subtypes for further study.

## Introduction

Infections with influenza A viruses cause short term respiratory infections with drastic health burdens and economic losses. Recurrent epidemics cause up to 650.000 deaths and 3 to 5 million cases of severe illness per year worldwide^1,2^. Two influenza subtypes are currently endemic in the human population: H3N2 influenza A (H3N2) viruses circulate since 1968 and H1N1 influenza A (pH1N1) viruses circulate since 2009^3-6^. Both lineages descend from pandemic outbreaks of reassortant viruses with segments of human and zoonotic origins^5^. The influenza genome consists of eight segments^3^ and encodes fourteen known proteins^7-10^. Reassortant viruses inherit viral segments from different viruses co-infecting the same cell, which, in case a novel HA is introduced, meet a largely naïve human population and thus can spread widely, causing a pandemic^3^. For H3N2 viruses, the polymerase basic protein 2 (PB1) and hemagglutinin (HA) segments are of avian origin, with evidence of adaptive processes, and the remaining ones originate from the previously circulating human H2N2 subtype^6,11^. The pH1N1 virus arose from a reassortment of three porcine viruses^5^, creating a virus with some segments of prior avian origins (NA, M; and another lineage with PB2 and PA), some distantly related to a human lineage circulating until 1998 (HA, NP, NS) and some descending more recently from humans (PB1)^5^.

Human influenza A viruses are the prime example of a rapidly evolving pathogen engaging in a co-evolutionary arms race with the host adaptive immune defenses. Due to their rapid evolution, the antigenic change of the major antigen to the humoral (B-cell) immune response, the surface protein HA, is monitored across the years^2,12-16^. Every couple of years, as susceptibility of the population becomes low due to prior infection or vaccinations, antigenically novel strains emerge and predominate in seasonal epidemics^17^. An antigenic match of vaccine viruses to the circulating viral population is achieved by regular updates of the vaccine composition^17,18^. Conversely, on the host side, upon infection or vaccination, specific antibodies are produced, recognizing B-cell epitopes (BCE) on HA, neuraminidase (NA) and matrix protein 2 (M2) and interfering with their function. In addition, cell-mediated immunity evokes T-cell responses against small peptides of internalized viral proteins, representing T-cell epitopes (TCE), which are digested and presented on the cell surface by the major histocompatibility complex (MHC). The innate immune system also recognizes pathogen-associated molecular patterns (PAMPs) as determinants of viruses and activates cellular antiviral responses, like activation of interferon (IFN)-induced proteins, to counteract viral replication^19,20^. In addition, human influenza viruses evolve towards resistance against antivirals^21^ and further adaptations to their host after initial establishment occur ^5^.

Evolutionary processes such as adaptation to specific environmental conditions can be studied on the level of genes, individual sites or even protein regions^15,16,22-29^. A widely used method determines the rates of non-synonymous to synonymous changes (*dN/dS* ratio) in a phylogeny^13,30^. A significant excess of *dN* to *dS*, or *dN/dS >* 1, provides evidence for positive selection, assuming that synonymous changes are neutral. This indicates that adaptation is taking place and that the changes of the respective genetic elements have led to a more favorable phenotype^31^. Methods measuring *dN / dS* should best be applied to analyze selection across, not within populations, and temporal influenza data may be considered as a series of independently evolving populations^32^.

While H3N2 viruses have been circulating for almost 50 years and presumably to do not require more adaptation to the human host, the recent introduction of pH1N1 viruses presents an excellent opportunity for a comparative analysis of the genome-wide evolutionary forces acting on both subtypes. We here sought to determine protein regions with signs of positive selection in both subtypes, and correlate them with known functional properties, to get a better understanding of the forces shaping the viral genome-wide evolution. We studied all available longitudinal sequence data for H3N2 and pH1N1 viruses collected since 1968 and 2009, respectively, across all eleven available protein structures. Though *dN/dS* values have been analysed already for sites^33^ or entire proteins of H3N2^34^ and pH1N1^35^, to our knowledge this is the first comparative and compressive study of the structures and functions linking to the genome-wide adaptation of both circulating influenza A subtypes.

## Methods

### Structure modeling

We analyzed the polymerase proteins PB2, PB1 and PA, the HA subunits 1 and 2, NP, NA and the splice-variants M1 and M2, as well as NS1 and NS2 for influenza subtypes H3N2 and pH1N1, respectively. Protein structures for HA for H3N2 viruses and NA for pH1N1 viruses were extracted from the RSCB database^36^. We modeled missing or incomplete protein structures with the homology modeling tool MODELLER using the A/Aichi/2/1968(H3N2) strain for H3N2 and the A/California/04/2009(H1N1) strain for pH1N1 as target sequences (Supplementary Table 1 & 2)^37^. The sequence identity between target sequence and the sequence of the homology models ranges from 67% (PB1) to 99% (HA1) (Supplementary Table 1 & 2). We were able to generate complete protein models for all polymerase subunits (both subtypes), HA and NS1 (both pH1N1) and partial protein models for the remaining proteins (Supplementary Table 1 & 2).

### Phylogenetic analysis

For each protein, we downloaded the amino acid sequences and the corresponding coding sequences from the NCBI flu database^38^. We removed identical coding sequences, which would accumulate on the same leaf in a phylogenetic tree because they contribute no additional information and removing significantly speeds up the calculation of alignments. The following steps were applied for each protein, individually:

We generated a multiple sequence alignment for amino acid sequences using muscle^39^. A subsequent removal of positions with more than 80% gaps using trimAL ensured a consistent numbering^40^. To speed up the analysis, we applied pal2nal to the protein alignment and the coding sequences to generate a multiple codon alignment^41^. Based on the multiple codon alignment, we inferred a phylogeny using FastTree with the GTR-model, which allows fast and precise inferences^42,43^. For each codon in the coding sequence and its corresponding amino acid, we reconstructed ancestral character states assigned to internal nodes using the Fitch algorithm with accelerated transformation (ACCTRAN) such that each internal node contains an intermediate sequence state of codon sequence^44^.

To measure positive selection, we calculate *dN/dS* ratios for every amino acid position considering each codon in the phylogenetic tree, separately^30,45^. To calculate the *dN*/*dS* statistics, we apply the tool described in Munch et al. (2017). Codon sequence changes are mapped onto branches in the phylogeny and are classified as synonymous or non-synonymous mutations. A synonymous mutation does not change the amino acid while a non-synonymous mutation does^13^. Starting from the root of the tree, we traverse down to the leaves and count position-wise every change that occurs between two nodes on the branches. For each site, we divide the amount of non-synonymous changes by the amount of synonymous changes normalized by the amount of comparisons to get the *dN/dS* ratio. To reduce the influence of sequencing artifacts, we exclude terminal branches and count changes supported by at least two viral strains.

### PatchDetection

The patch detection method is based on the methods from Tusche et al. (2012) and Kratsch et al. (2016). It includes a graph cut algorithm combining spatial information of structure models with *dN/dS* measurements. Following the original versions, we create a graph with nodes *n* representing protein sites that are connected to all neighboring residues within a radius of *δ*. The edges are weighted by a distance function *e*^−*dist*(*m*, *n*)^ were *dist*(*m*, *n*) calculates the euclidean distance between the *C_α_*-atoms of a residue pair *n* and *m*. Nodes with a close spatial proximity therefore have a strong connection. We add two additional nodes to the graph, the positive selection node (*pos*) and the negative selection node (*neg*). Each *n* is connected to *pos* with *P*(*n*) and to *neg* with 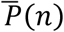. Sites are separated into a positive or a negative selection set by a minimum graph cut approach. The cut divides the graph into two halves by minimizing the sum *C* of weights from edges connecting these halves with a cost function:

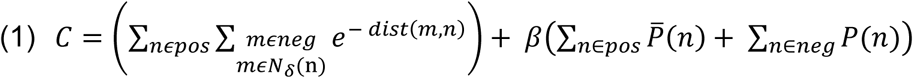

In a final step, sites in the positive selection set are merged into patches together with neighboring sites within a distance of *δ*.

In the new formulation, we adjusted the graph-cut function to directly include *dN/dS* values.. We set *P*(*n*) = *dN/dS* and 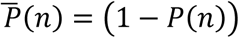 to weigh the positive selection node (*pos*) and the negative selection node (*neg*), balancing positively-selected sites against sites not under positive selection. We applied a radius in the stable interval between 7 and 8.5 Å (see approach Supplementary Table 2 in Kratsch et al. (2016)). We include buried and exposed sites to account for macromolecular changes in the core of the protein. To determine *β*, we start at 1, incrementally increase *β* and calculate patches at each step until patches do not change anymore within 100 steps. Having reached saturation at *β^max^*, we set *β* = *β^max^* * 0.5 to balance between both extreme cases in which either the distance function dominates at *β* = 0 or the *dN/dS* term dominates at *β^max^*.

We evaluate the robustness of each patch by subsampling protein sites. This provides statistical support to estimate the stability of each patch regarding its site content. We define a set of patches for a protein as *P =* {*p*_1_, *p*_2_,..,*p_n_*}, where *n* is the total number of patches and *p_i_* a set of sites under positive selection in the *i*-th patch. We generate *N* = 1.000 samples by randomly removing 10% of sites from the protein structure and define *P_j_* = {*p_j_*_,1,_ *p_j,_*_2_,*..,p_j,k_*} as the *k* patches that we re-calculate on the *j*-th sample. For each patch *p_i_* ∈ *P*, we perform the following steps: For each sample *j*, we define the patch *p_max_* ∈ *P_j_*, which has the maximum number of overlapping sites with *p_i_*, and calculate the number of true positives as 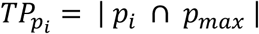, false negatives as 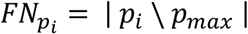 and false positives as 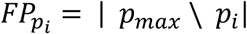. Sites that were removed by the sampling process are omitted for calculation of *TP_i_, FN_i_, and FP_i_*. We use the standard definitions for recall, precision and F-score^46^. The criterion 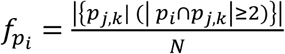 reflects how stable a patch appears in the samples. When the value approaches zero, the patch tends to disappear, a value of one indicates that the patch is stable and a value larger than one indicates an unstable patch that breaks apart in the subsamples. In addition, we provide the average *dN/dS* value per patch.

### NA activity assays

The NA activity of pH1N1 influenza A virus (A/Hamburg/NY1580/2009) was measured by a fluorescence-based assay using the fluorogenic substrate 2’-(4-Methylumbelliferyl)-α-D-N-acetylneuraminic acid (4-MU-NANA, Sigma-Aldrich) which is cleaved by the viral NA enzyme to release the fluorescent product 4-methylumbelliferone (4-MU)^47^. Therefore, recombinant wildtype (wt) or NA mutant (D199N, D199E, S247N, S247G) pH1N1 viruses were diluted to an equivalent NA activity and pre-incubated with 10, 100 or 1000 nM Oseltamivir (Tamiflu, Roche) or mock treated with 1xPBS for 15 min at 37°C. The NA enzyme activity was initiated by adding μM 4-MU-NANA in calcium-TBS (6.8 mM CaCl_2_, 0.85 % NaCl, 0.02 M Tris; pH7.3). After 30 min at 37°C, the reaction was stopped by the addition of 100 μl of 0,1 M glycine buffer (pH 10,7) containing 25 % ethanol. The fluorescence of released 4-MU was determined with a Safire 2 multi-plate reader (Tecan) using excitation and emission wavelengths of 355 nm and 460 nm, respectively. The mean fluorescence signal of substrate without virus was subtracted as background from the signals obtained in the other wells. The specific NA activity was expressed in percentage.

## Results

### Global trends

We calculated *dN/dS* values for all proteins of pH1N1 and H3N2 viruses (Supplementary Figure 1). Notably, the HA1 subunit, historically considered the main driver of adaptation of seasonal influenza viruses^12,13^, did not show the most evidence for positive selection. It was ranked fifth for both subtypes when jointly analyzing all time periods, as indicated in analyses on smaller data sets ^34,35^. Instead, NA had the highest mean *dN/dS* value for both subtypes, followed by M2, NS1 and NS2. NP and the PB1 proteins had the lowest mean *dN/dS* value in both subtypes (Supplementary Table 3). We confirmed this finding by re-calculating the *dN/dS* statistics using the Suzuki-Gojobori counting approach implemented in HyPhy SLAC^48^ (Supplementary Table 3).

Plotting the mean protein *dN*/*dS* values over time shows this trend and highlights individual seasons in which the mean *dN*/*dS* peak (Supplementary Figure 2). Contrary to mean *dN*/*dS* values for the entire time period, in a season-wise analysis, the mean *dN*/*dS* of HA1 surpassed the other proteins for both subtypes, where a tendency of higher values in winter than in summer seasons correlated with more available sequences (Supplementary Figure 2). This indicates that across protein comparisons in early studies or a season-wise analysis of the current data may have limited value due to a lack of available data for most proteins and also due to the nature of the *dN*/*dS* statistic^32^.

The Kolmogorov–Smirnov-test (KS-test; *H*_0_: *dN*/*dS* distribution of pH1N1 protein is smaller than *dN*/*dS* distribution of H3N2 protein) showed that the global *dN*/*dS* distributions of pH1N1 proteins were significantly larger than those of H3N2 proteins, except for HA2, M2 and NS1 (Supplementary Table 4). This was especially apparent for the polymerase proteins PB2, PB1 and PA, which are of known importance for host adaptation^49^. We hypothesize that positive selection was acting on substantially more regions for pH1N1 than for H3N2 viruses recently, due to their requirement to further adapt to the new host population, which the H3N2 virus likely has undertaken in the early years of its circulation since the 1968 pandemic, for which little data is available.

Following the methodology in Klingen et al. (2018), we calculated the average number of sweep-related amino acid changes for pH1N1 and H3N2 viruses and compare the number of sweeps fixed over time across all eight segments (Supplementary Figure 3). The number of sweep-related changes becoming fixed per year was greatest for HA1 and NA in both viruses, but substantially increased in H3N2 relative to pH1N1 viruses, in line with a lesser need for antigenic immune escape for the latter in the early years of circulating in the human population. For M2 and NS2, it was the opposite case and all remaining proteins had similar rates. Taken together, positive selection has the strongest effect in both viruses on the major surface proteins, demonstrates by their high rates of selective sweeps, while the nature of the segments suboptimally adapted with positive selection acting on them differ substantially between them.

To investigate this further, we identified clusters (patches) of sites under positive selection on the protein structure, based on *dN*/*dS* values and structure models, using a methodology based on our prior work described in Tusche et al. (2012) and Kratsch et al. (2016) (Methods). This was done for all proteins in both subtypes for which we have a suitable homology model and sufficient evidence for positive selection within the regions of the available partial structures (Table 1). The number of patches and of sites clustered into patches for pH1N1 viruses exceeded those for H3N2 viruses, indicating that more regions of pH1N1 are currently suboptimally adapted to the human host. This was despite of H3N2 data covering a longer time span (1968-2016; though data largely originates from after 1999) than for pH1N1 (2009-2016). Specifically, for the polymerase subunits PB2 (17 vs. 6), PB1 (7 vs. 4) and PA (12 vs. 7), pH1N1 viruses had more patches than H3N2, as well as for NP (7 vs. 1) and NA (15 vs. 8). There were fewer differences for NS1 (11 vs. 9 patches) and M2 (3 vs. 2), indicating a similar degree of positive selection acting on NS1 and M2 for both subtypes. We inferred two patches for pH1N1/M1 and no patches for H3N2/M1 and for NS2 (both subtypes). Only for the major surface antigen HA we detected fewer patches in pH1N1 viruses than in H3N2 viruses (12 vs. 13 for HA1 and 0 vs. 1 for HA2), in agreement with pH1N1 experiencing less pressure to change antigenically and escape accumulating immunity in the host population over the past decade than H3N2 viruses (see below).

**Table 1:**
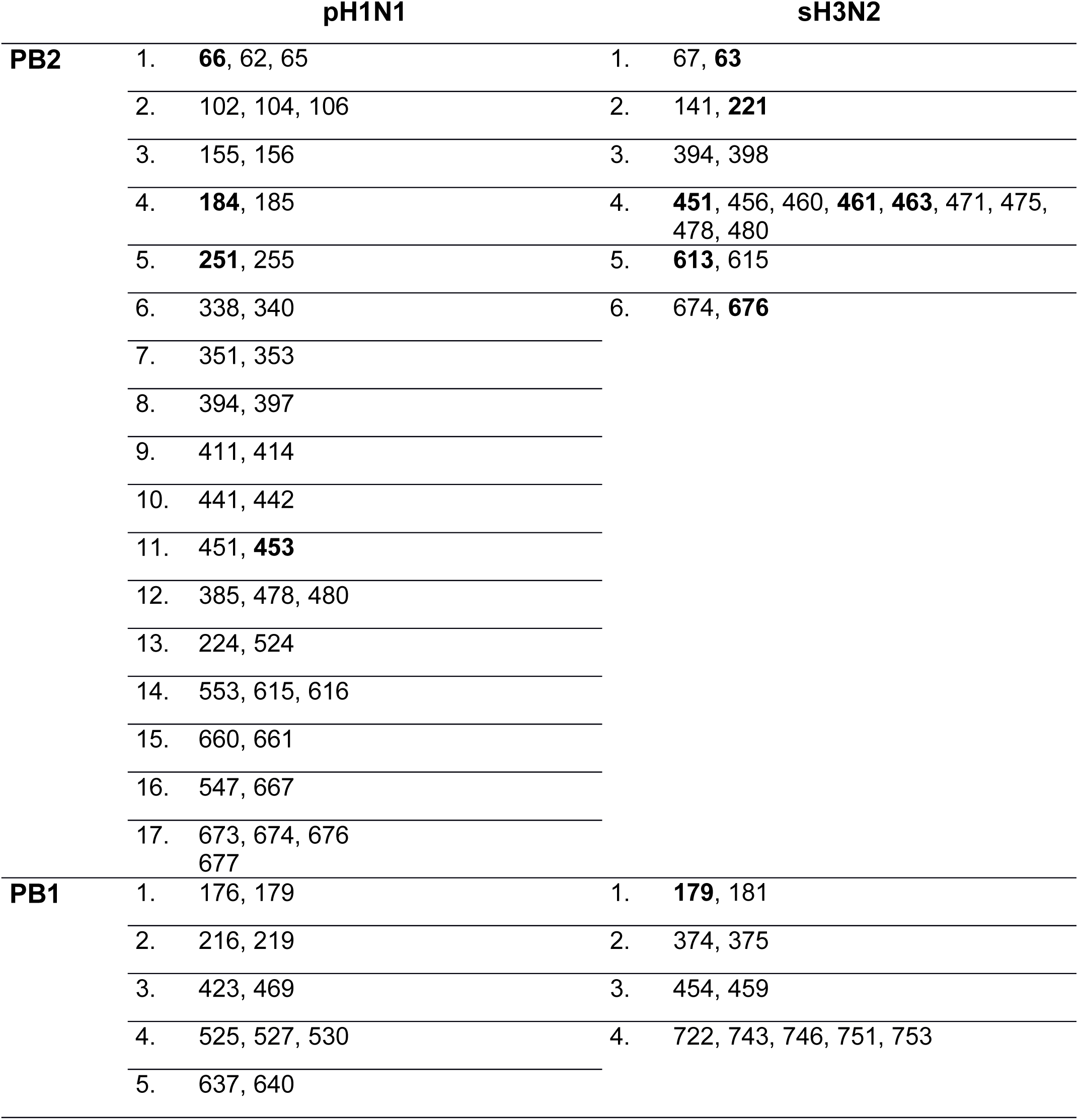
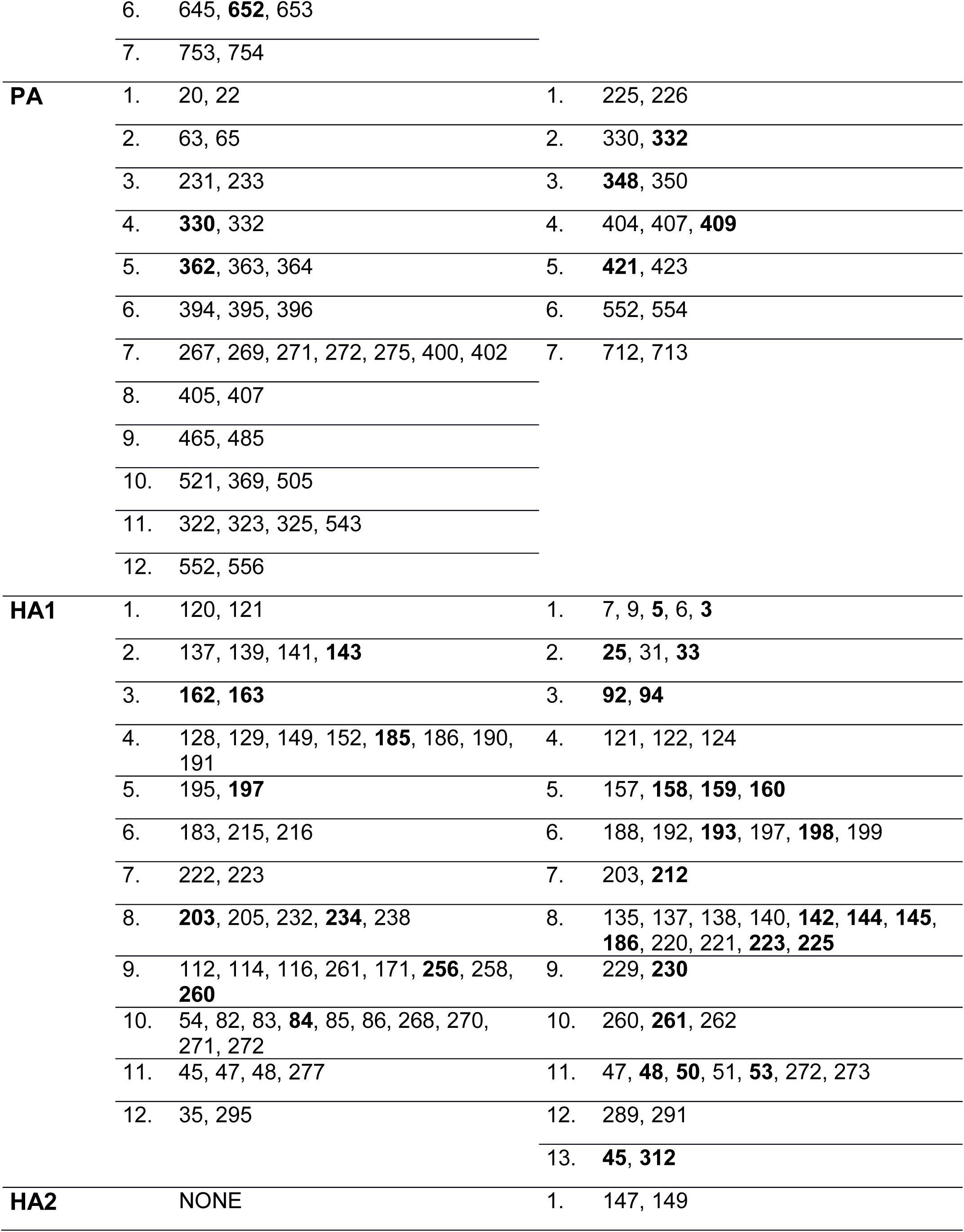
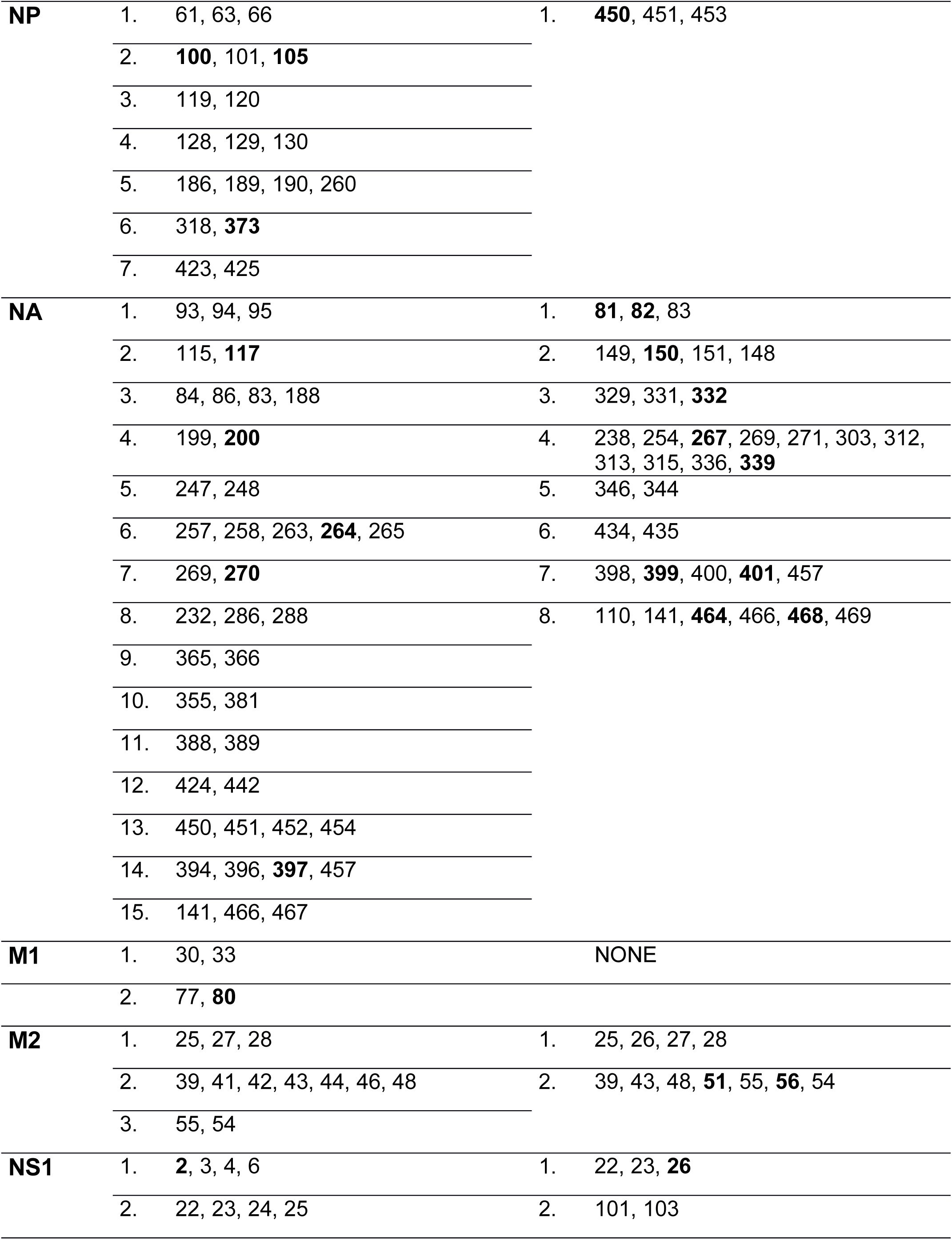
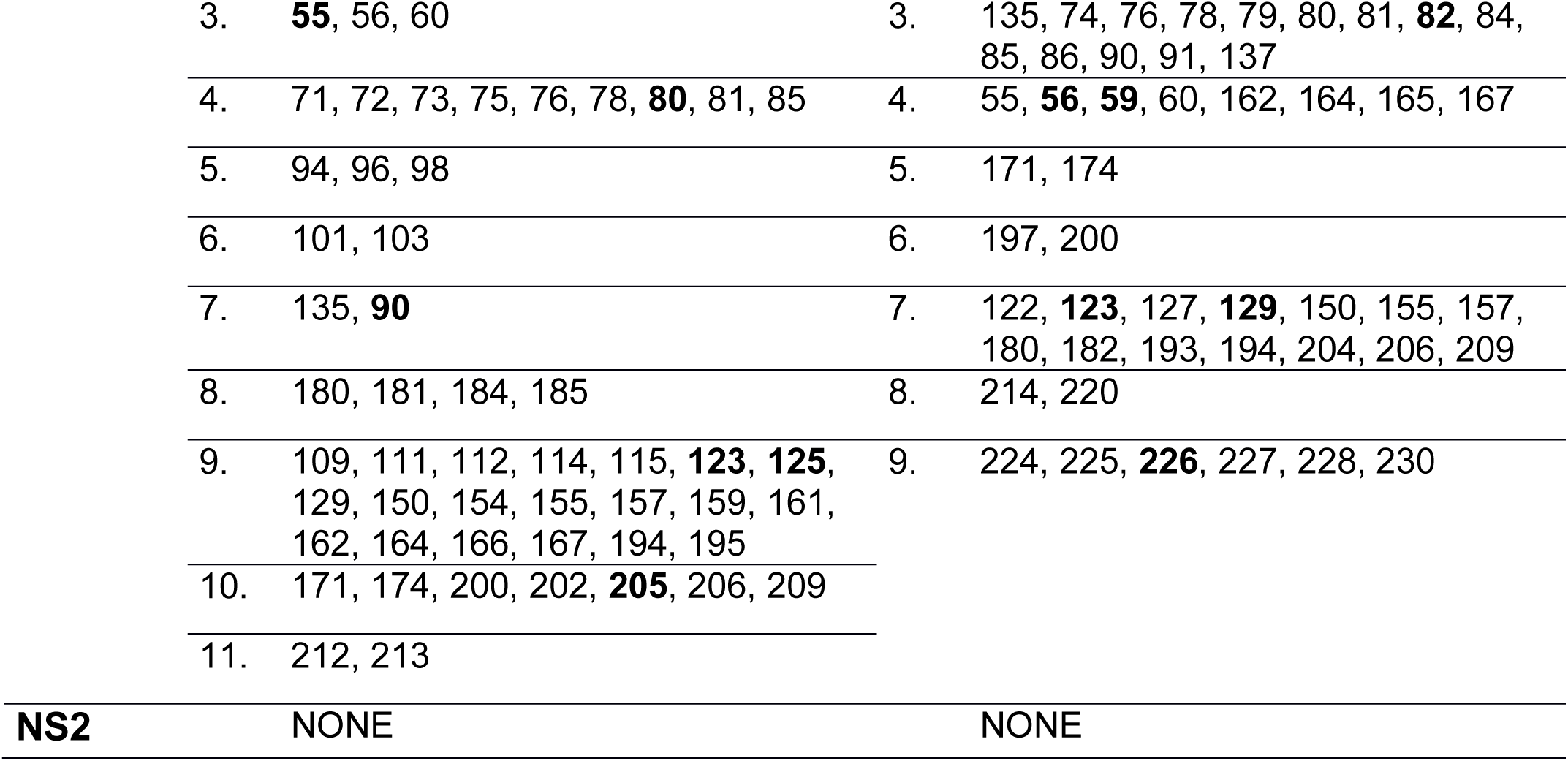
Patches and residues under positive selection for all analyzed influenza proteins. Overview of all patches and patch sites for each protein of pH1N1 and H2N3 viruses. Sweep-related sites are marked in bold.

|  | pH1N1 | sH3N2 |
| --- | --- | --- |
| **PB2** | 1. **66**, 62, 65 | 1. 67, **63** |
|  | 2. 102, 104, 106 | 2. 141, **221** |
|  | 3. 155, 156 | 3. 394, 398 |
|  | 4. **184**, 185 | 4. **451**, 456, 460, **461**, **463**, 471, 475, 478, 480 |
|  | 5. **251**, 255 | 5. **613**, 615 |
|  | 6. 338, 340 | 6. 674, **676** |
|  | 7. 351, 353 |  |
|  | 8. 394, 397 |  |
|  | 9. 411, 414 |  |
|  | 10. 441, 442 |  |
|  | 11. 451, **453** |  |
|  | 12. 385, 478, 480 |  |
|  | 13. 224, 524 |  |
|  | 14. 553, 615, 616 |  |
|  | 15. 660, 661 |  |
|  | 16. 547, 667 |  |
|  | 17. 673, 674, 676<br>677 |  |
| **PB1** | 1. 176, 179 | 1. **179**, 181 |
|  | 2. 216, 219 | 2. 374, 375 |
|  | 3. 423, 469 | 3. 454, 459 |
|  | 4. 525, 527, 530 | 4. 722, 743, 746, 751, 753 |
|  | 5. 637, 640 |  |

|  |  |  |  |  |
| --- | --- | --- | --- | --- |
|  | 6. | 645, **652**, 653 |  |  |
|  | 7. | 753, 754 |  |  |
| **PA** | 1. | 20, 22 | 1. | 225, 226 |
|  | 2. | 63, 65 | 2. | 330, **332** |
|  | 3. | 231, 233 | 3. | **348**, 350 |
|  | 4. | **330**, 332 | 4. | 404, 407, **409** |
|  | 5. | **362**, 363, 364 | 5. | **421**, 423 |
|  | 6. | 394, 395, 396 | 6. | 552, 554 |
|  | 7. | 267, 269, 271, 272, 275, 400, 402 | 7. | 712, 713 |
|  | 8. | 405, 407 |  |  |
|  | 9. | 465, 485 |  |  |
|  | 10. | 521, 369, 505 |  |  |
|  | 11. | 322, 323, 325, 543 |  |  |
|  | 12. | 552, 556 |  |  |
| **HA1** | 1. | 120, 121 | 1. | 7, 9, **5**, 6, **3** |
|  | 2. | 137, 139, 141, **143** | 2. | **25**, 31, **33** |
|  | 3. | **162**, **163** | 3. | **92**, **94** |
|  | 4. | 128, 129, 149, 152, **185**, 186, 190, 191 | 4. | 121, 122, 124 |
|  | 5. | 195, **197** | 5. | 157, **158**, **159**, **160** |
|  | 6. | 183, 215, 216 | 6. | 188, 192, **193**, 197, **198**, 199 |
|  | 7. | 222, 223 | 7. | 203, **212** |
|  | 8. | **203**, 205, 232, **234**, 238 | 8. | 135, 137, 138, 140, **142**, **144**, **145**, **186**, 220, 221, **223**, **225** |
|  | 9. | 112, 114, 116, 261, 171, **256**, 258, **260** | 9. | 229, **230** |
|  | 10. | 54, 82, 83, **84**, 85, 86, 268, 270, 271, 272 | 10. | 260, **261**, 262 |
|  | 11. | 45, 47, 48, 277 | 11. | 47, **48**, **50**, 51, **53**, 272, 273 |
|  | 12. | 35, 295 | 12. | 289, 291 |
|  |  |  | 13. | **45**, **312** |
| **HA2** |  | NONE | 1. | 147, 149 |
| **NP** | 1. | 61, 63, 66 | 1. | **450**, 451, 453 |
|  | 2. | **100**, 101, **105** |  |  |
|  | 3. | 119, 120 |  |  |
|  | 4. | 128, 129, 130 |  |  |
|  | 5. | 186, 189, 190, 260 |  |  |
|  | 6. | 318, **373** |  |  |
|  | 7. | 423, 425 |  |  |
| **NA** | 1. | 93, 94, 95 | 1. | **81**, **82**, 83 |
|  | 2. | 115, **117** | 2. | 149, **150**, 151, 148 |
|  | 3. | 84, 86, 83, 188 | 3. | 329, 331, **332** |
|  | 4. | 199, **200** | 4. | 238, 254, **267**, 269, 271, 303, 312, 313, 315, 336, **339** |
|  | 5. | 247, 248 | 5. | 346, 344 |
|  | 6. | 257, 258, 263, **264**, 265 | 6. | 434, 435 |
|  | 7. | 269, **270** | 7. | 398, **399**, 400, **401**, 457 |
|  | 8. | 232, 286, 288 | 8. | 110, 141, **464**, 466, **468**, 469 |
|  | 9. | 365, 366 |  |  |
|  | 10. | 355, 381 |  |  |
|  | 11. | 388, 389 |  |  |
|  | 12. | 424, 442 |  |  |
|  | 13. | 450, 451, 452, 454 |  |  |
|  | 14. | 394, 396, **397**, 457 |  |  |
|  | 15. | 141, 466, 467 |  |  |
| **M1** | 1. | 30, 33 |  | NONE |
|  | 2. | 77, **80** |  |  |
| **M2** | 1. | 25, 27, 28 | 1. | 25, 26, 27, 28 |
|  | 2. | 39, 41, 42, 43, 44, 46, 48 | 2. | 39, 43, 48, **51**, 55, **56**, 54 |
|  | 3. | 55, 54 |  |  |
| **NS1** | 1. | **2**, 3, 4, 6 | 1. | 22, 23, **26** |
|  | 2. | 22, 23, 24, 25 | 2. | 101, 103 |

|  |  |  |  |
| --- | --- | --- | --- |
| 3. | **55**, 56, 60 | 3. | 135, 74, 76, 78, 79, 80, 81, **82**, 84,<br>85, 86, 90, 91, 137 |
| 4. | 71, 72, 73, 75, 76, 78, **80**, 81, 85 | 4. | 55, **56**, **59**, 60, 162, 164, 165, 167 |
| 5. | 94, 96, 98 | 5. | 171, 174 |
| 6. | 101, 103 | 6. | 197, 200 |
| 7. | 135, **90** | 7. | 122, **123**, 127, **129**, 150, 155, 157,<br>180, 182, 193, 194, 204, 206, 209 |
| 8. | 180, 181, 184, 185 | 8. | 214, 220 |
| 9. | 109, 111, 112, 114, 115, **123**, **125**,<br>129, 150, 154, 155, 157, 159, 161,<br>162, 164, 166, 167, 194, 195 | 9. | 224, 225, **226**, 227, 228, 230 |
| 10. | 171, 174, 200, 202, **205**, 206, 209 |  |  |
| 11. | 212, 213 |  |  |
| **NS2** | NONE |  | NONE |

### Adaptive immune evolution

We investigated which patches are located in regions that act as BCE or TCE and thus play a role in evading the host’s immune response. In addition to extracellular regions of HA and NA, the M2 ion channel can be partly recognized as a BCE, too^33^. TCE epitopes exist in intracellular regions of HA, NA and M2^33^.

Of the H3N2/HA1 sites, 15% percent (52 sites) are located in a patch. Antigenic or BCE definitions for H3N2/HA1 were originally proposed by Wiley et al. (1981) and Wiley and Skehel (1987). Subsequent studies refined key regions or sites responsible for antigenic drift in HA1^14,23,52,53^. Thus, we also compared H3N2 patch sites with “antigenic patch” sites^23^, “adapatch” sites^22^, key antigenic sites^52^, sweep-related sites^16^ and TCE sites^33^ (Figure 1A). Eight of twenty-three antigenic patch sites are included in patches (34.7%). Of the seven key antigenic sites by Koel et al. (2013), sites 155, 156 and 189 were excluded from a patch. These findings underline that antigenic alterations cannot exclusively explain positive selection^14,16^. 62.8% of adapatch sites overlap with the patch sites of this study. In turn, thirty (57.7%) sites were newly clustered into patches, mostly due to inclusion of both exposed and, newly, also buried protein sites in patches. Half of the sweep-related sites from ^16^ match patch sites. While sweep-related sites are likely under positive selection and may become fixed in the circulating viral population, they do not necessarily have large *dN*/*dS* values, which would require multiple changes at a given site for detection. Of patches clustering on the globular head of HA1, patches 6, 8 and 9 are vicinal to receptor binding sites (RBS) 98, 153 and 195 ^54^ (Figure 2A). Altogether, the overlap of patch sites with BCE sites was ~ 70% and of patch sites with TCE sites was only ~20%, in line with evasion of humoral immune responses being more pronounced for the viral major surface protein than escape from cell-mediated immune evasion.

**Figure 1:**
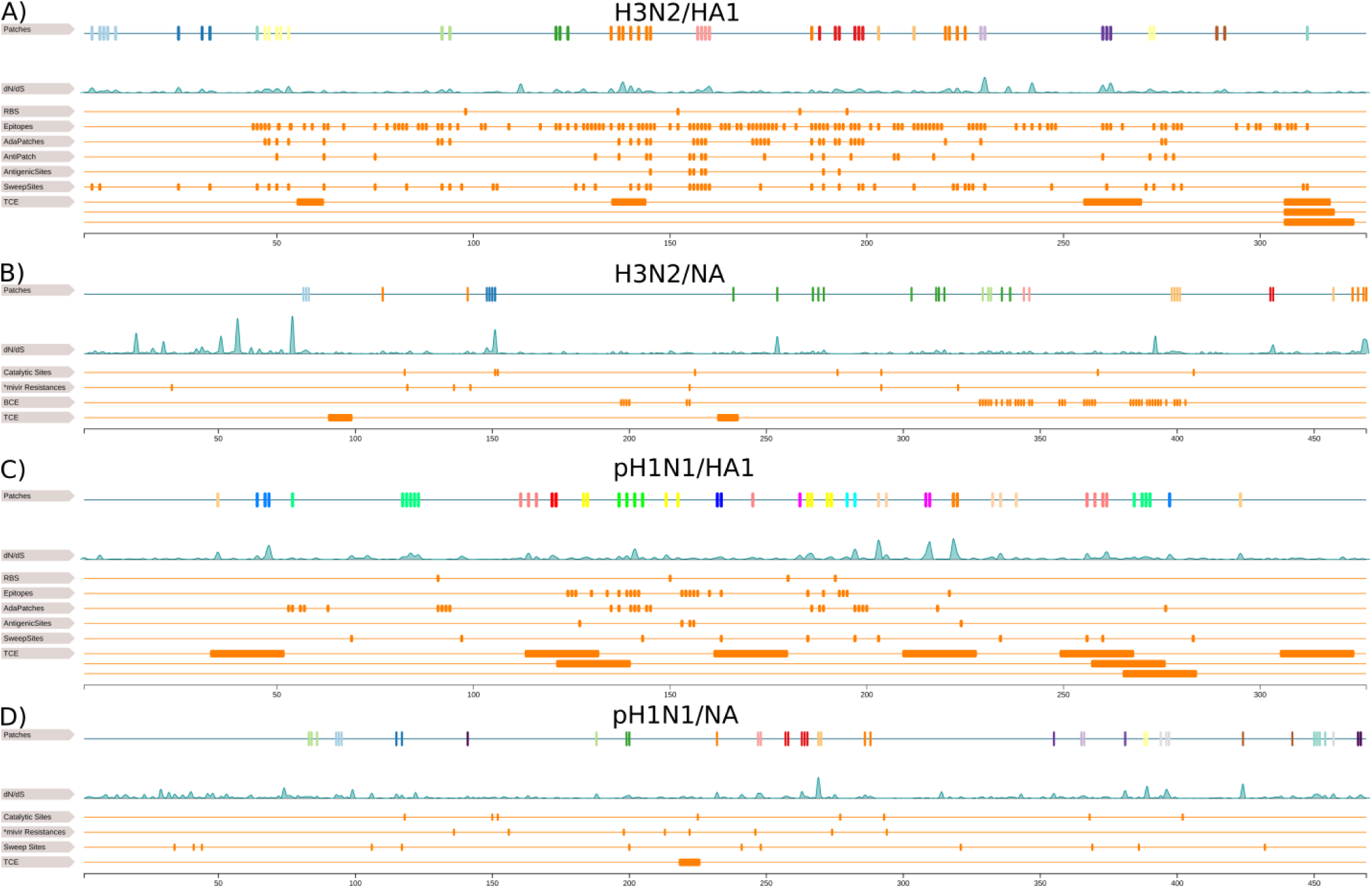
Linear representation of surface proteins. This figure illustrates the overlap of patch sites with known functional sites in H3N2/HA1 **(A)** H3N2/NA **(B)** pH1N1/HA1 **(C)** and pH1N1/NA **(D)**. In each subfigure, the first row is a linear representation of patches and the second row shows the *dN/dS* distribution of the protein. In **(A)**, we compare with the RBS^54^, BCE sites^50,51^, “adapatch” sites^22^, “antigenic patch” sites^23^, key antigenic sites^52^, sweep-related sites^16^ and TCE sites^33^. In **(B)**, we compare with the catalytic sites^75^, oseltamivir and zanamivir resistance sites^21^, BCE sites and TCE sites^33^. In **(C)**, we compare with the RBS^59^, BCE^55,56^, “adapatch” sites^22^, key antigenic sites^57^, sweep-related sites^16^, TCE sites^33^. In **(D)**, we compare with catalytic sites^75,76^, oseltamivir and zanamivir resistance sites^21^, sweep-related sites^16^ and TCE sites.

Epitopes in HA1 for pH1N1 viruses were originally defined for seasonal H1N1 viruses by Caton et al. (1982). As the pH1N1 replaced the seasonal H1N1 virus, Matsuzaki et al. (2014) redefined the former epitopes by mapping them onto the pH1N1/HA1 protein. We match patches in pH1N1/HA1 with “adapatch” sites^22^, key antigenic sites^57^, sweep-related sites^16^, TCE sites^58^ and BCE sites^56^ (Figure 1C). In total, 16% percent of pH1N1/HA1 sites are located in patches. Fifteen of twenty-five “adapatch” sites overlap with novel patch sites (60%) and forty of fifty-five novel patch sites are no adapatch sites (70%). In comparison to H3N2/HA, this is a larger discrepancy that cannot only be explained by the inclusion of buried sites. A second reason is the longer time period for which we could include pH1N1 virus in this study. 81% of sweep-related sites are also patch sites, which is a larger overlap compared to H3N2/HA1 data. Koel et al. (2015) described five key antigenic positions (127, 153, 155, 156 and 224) for sH1N1/HA1, of which none is located in a patch. However, most of them are in the vicinity of patch 4, which also surrounds RBS 150 and 192^59^ (Figure 2B). For pH1N1/HA1 we observe almost identical overlaps of patch sites with TCE sites (~20%) and BCE sites (~26%). The low percentage of TCE sites in patches for pH1N1/HA1 is similar to H3N2/HA1 results, but, the overlap with BCE sites is lower compared to H3N2 data. This likely is also indicative of pH1N1 viruses not having to evade pre-existing immunity yet as much as H3N2 viruses and that most positive selection observed for HA does not result from humoral immune evasion.

**Figure 2:**
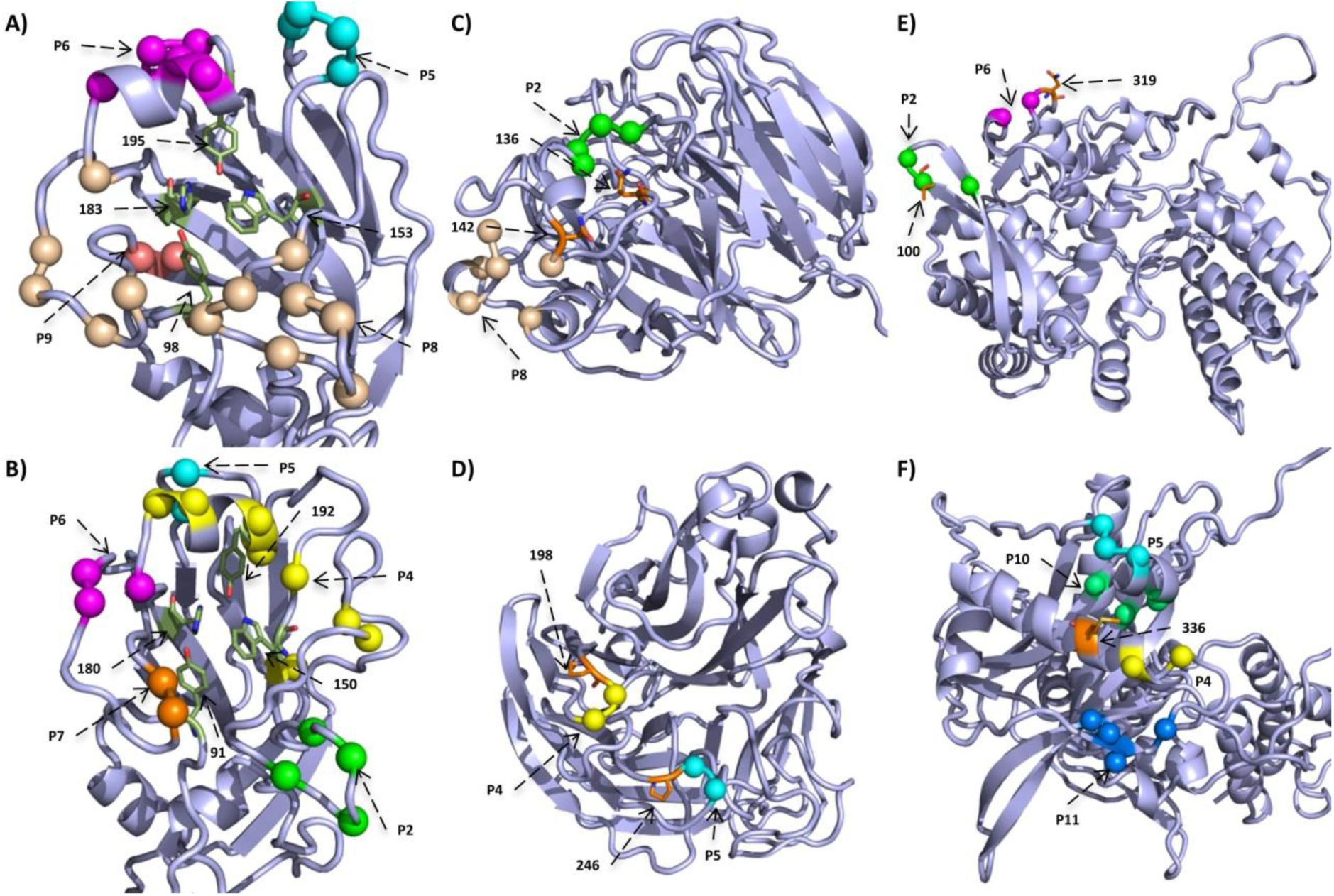
Structural representation of selected patches. We mapped selected patches on the corresponding protein models shown as cartoon in light blue with interesting residues shown as sticks and colored by atom-type (green and orange for carbon, red for oxygen and blue for nitrogen). The structures in **(A)** and **(B)** depict the head region of H3N2/HA1 and pH1N1/HA1, respectively, and show the RBS^54,59^ and surrounding patches. In **(C)** and **(D)**, we show views on the H3N2/NA and pH1N1/NA, respectively, and highlight patches that likely extend resistance sites (orange). In pH1N1/NP patch 2 and 6 are close to known mammalian adaptation sites 100^16,80,81^ and site 319 (orange)^82^, respectively **(E)**. The structure of pH1N1/PA is shown in **(F)** and highlights adaptation site 336 (orange) that increases polymerase activity in mammalian cells and neighboring patches 4, 5, 10 and 11.

Patch sites of H3N2/NA match 27% reported BCE sites^60^. While the overlap of patch sites with TCE for H3N2/NA is ~2%, there is no overlap for pH1N1/NA with known TCE sites, indicating that there was little immune evasion of TCR of NA (Figure 1B&D). We could not investigate for H3N2/M2 and overlap of BCE and TCE sites with patch sites, because the former are located in regions not covered by our homology model (Supplementary Table 2).

### Attenuation of innate immune responses

NS1 suppresses multiple antiviral host responses and mediates viral replication. Interferon synthesis is suppressed by binding the cleavage and polyadenylation specificity factor (CPSF30) and interfering with IFN-β pre-mRNA synthesis^61^. 22.3% (Fifty-three sites) of protein sites in H3N2 viruses and 27.4% (sixty sites) of protein sites of pH1N1 viruses in NS1 are clustered in a patch, which is the largest number for all proteins (Supplementary Table 5). In both subtypes, most patches occur in the C-terminal effector domain (85-end of protein), while the N-terminal RNA-binding domain (1-73) is almost conserved across subtypes^62^. A flexible linker region of 12 conserved residues connects both distinct domains ^63^. In both subtypes, a large patch spans the linker region, but excludes sites 69 and 77 that are critical to maintain dimerization^64^. In H3N2 viruses, changes at sites 103 and 106^65^ and in the binding pocket (110, 117, 119, 121, 180, 183, 184 and 187), which interacts with the second and third zinc finger domain of CPSF30 ^66^, link to post-transcriptional inhibition of antiviral IFN-stimulation^61^. Of these sites, patch 2 includes site 103 and patch 7 is next to sites 119, 121 and 183, and includes site 180 (Figure 3C). In addition, the sites 108, 125 and 189 are relevant for CPSF30 binding in pH1N1 viruses^65,66^. In pH1N1, they overlap with patch 6 (site 103), patch 9 (site 125) and are adjacent to patch 9 (sites 106 and 108) (Figure 3D). Clark et al. (2017) showed that substitutions at sites 55, 90, 123, 125, 131 and 205 inhibit host gene expression primarily by inhibiting CPSF30^67^, representing an adaptation of pH1N1 to humans^16,67^. Site 125 was also described as a host adaptation site in Selman et al. (2012). Of these sites, site 55 is part of patch 3, site 90 of patch 7, site 123 and 125 of patch 9 and 205 of patch 10 (Figure 3D). NS1 also interferes with the expression of transcription factor IRF3 and NF-κB, which initiate IFN transcription^69^. The suppression of IFN responses is mediated via site 196 that blocks the IFN cascade when the amino acid E is present^69^. Site 196 is proximal to patch 6 in H3N2 and patch 9 in pH1N1. In H3N2 viruses, sites 189 and 194 affect interferon responses^70^ and in pH1N1 viruses, changes in site 171 reduce host gene expression^62^. Site 194 is included in patch 7, while the pH1N1 specific site 171 lies in patch 10. A different viral strategy is directly binding of NS1 region 123-127 to protein kinase PKR, suppressing its activation and downstream antiviral responses^61,71,72^. This region overlaps with patch 9 in pH1N1/NS1 and patch 7 in H3N2/NS1 (Figure 3A&B). Thus, patch 9 in pH1N1/NS1, which has a high average *dN/dS* value (~1.96), and patch 7 in H3N2 /NS1, with a high average *dN/dS* value (~1.2), may be relevant for antagonizing IFN responses via CPSF30-binding and suppression of protein kinase PKR in both subtypes. Taken together, our findings indicate that in both subtypes, changes that increase the suppression of antiviral host responses are occurring, and that a substantial part of the NS1 protein is relevant for this process. Strikingly, this is also the case for H3N2, despite of its long-term circulation in the human population. Patches in the N-terminal RNA-binding domain do not overlap with functionally relevant sites such as dsRNA-binding sites (35, 37, 38 and 41)^7,73^ and sites that inhibit IFN induced 2’, 5’-Oligoadenylate Synthetase (OAS) (38 and 41), which both prevent NF-κB activation and IFN-β induction^61,73^.

**Figure 3:**
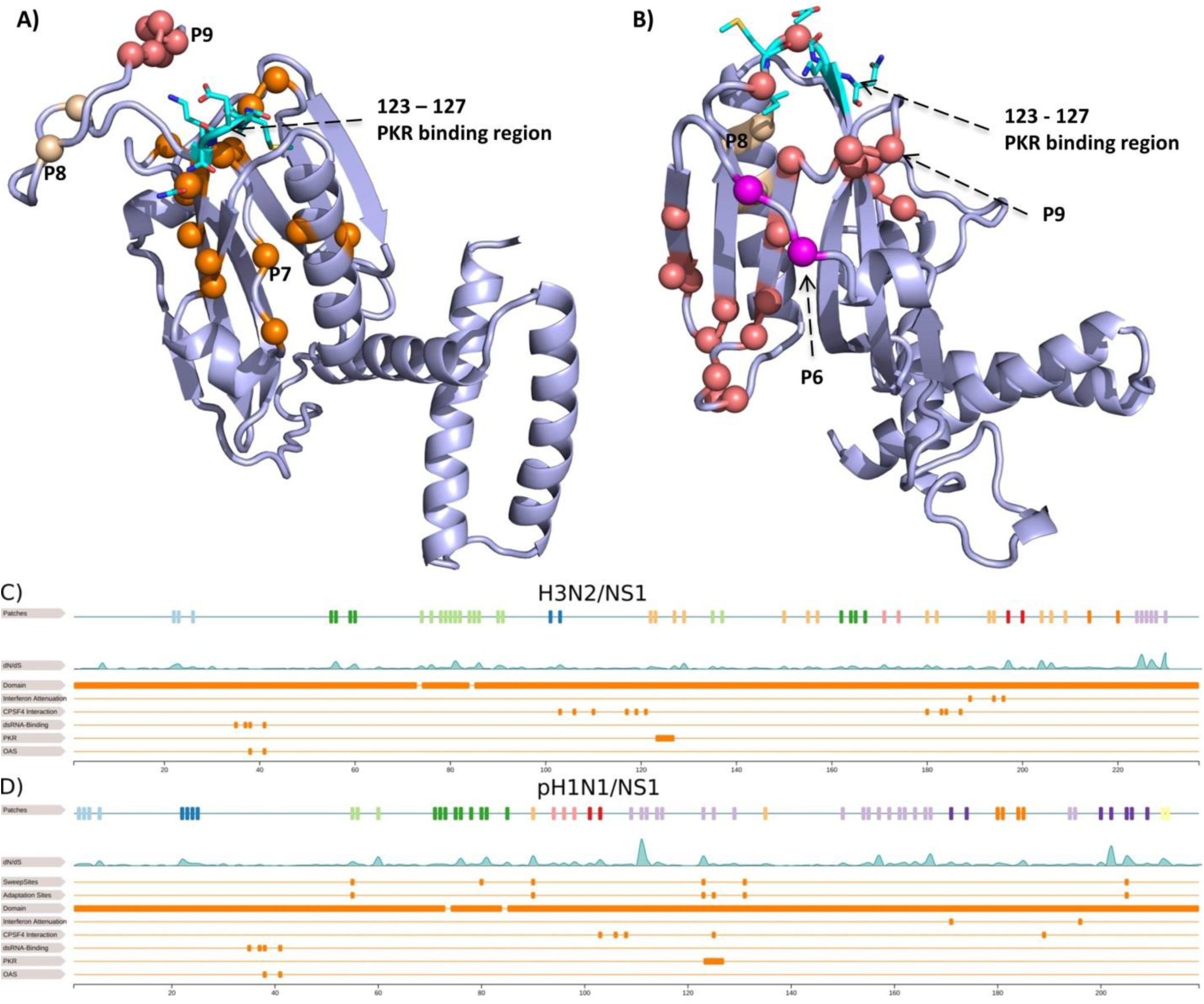
Structural and linear representation of NS1 patches. The NS1 models for H3N2 in **(A)** and pH1N1 in **(B)** are shown as cartoon in light blue with the PKR binding region^61,71,72^ shown as sticks and colored by atom-type (cyan for carbon, red for oxygen and blue for nitrogen). We highlight patches in the surrounding region of the PKR binding region with a major focus on patch 7 in H3N2 and patch 9 in pH1N1. In **(C)** and **(D)**, the first row is a linear representation of patches and the second row shows the distribution of H3N2/NS1 and pH1N1/NS1. The third and fourth row in **(D)** show sweep-related sites^16^ and adaptations sites^67^ in pH1N1/NS1. Remaining rows in both subfigures depict the protein domains^62,63^, interferon attenuation sites^62,69,70^, CPSF30 interaction sites^61,65,66^, dsRNA binding sites^7,73^, PKR^61,71,72^ and OAS binding sites^61,73^.

### Resistance evolution

The ion channel of M2 is targeted by the antivirals amantadine and rimantadine, which are used to treat influenza infections^21^. We found a strong link between patch sites and antiviral resistances in the M2 protein. Known amantadine resistance sites 26, 27, 30, 31, 34 and 38, are surrounded by or part (site 27) of patch 1 and are also in vicinity of patch 2 in pH1N1^21^. In H3N2, patch 1 includes amantadine resistance sites (26 and 27). Rimantadine resistance sites 40, 41, 42 and 44 are included or in the vicinity of patch 2 in pH1N1 and H3N2. Thus, there are similar trends for pH1N1 and H3N2 patches in M2, and all patch sites are either part of or next to known resistance sites, indicating their relevance for resistance development.

Currently, both drugs are not recommended anymore, because of increasing resistances in circulating viruses^74^. Alternatives are oseltamivir and zanamivir that inhibit NA sialidase, preventing viral detachment from the host cell^21^. In both subtypes, two patches are linked to known antiviral resistance sites, providing evidence for positive selection against these two drugs in NA for seasonal influenza viruses (2 out of 8 in H3N2 and 2 out of 15 in pH1N1): resistances against the anti-NA drugs are known to be linked to sites in NA 33, 119, 136, 142, 292, 320 and 222 in H3N2 viruses and 136, 156, 198, 213, 222, 246, 274 and 294 in pH1N1 viruses^21^. In H3N2/NA, resistance site 136 is in vicinity of patch 2 and 142 is next to patch 8 (Figure 2C). Resistance sites 198 and 246 are next to patch 4 and patch 5 in pH1N1/NA, respectively (Figure 2D). In order to assess whether the predicted sites confer resistance against NA inhibitors, such as oseltamivir, we experimentally investigated positions 199 and site 247 of patch 4 and patch 5, respectively, for a role in generating oseltamivir resistances in NA activity assays. We show that amino acid substitutions in NA from D to N at position 199 and from S to N at position 247 considerably reduced the inhibitory effect of oseltamivir up to 30% neuraminidase activity upon treatment with 100nm oseltamivir (Figure 4). The ability of these mutations to confer oseltamivir resistance was further increased upon treatment with 1000nm oseltamivir concentrations, further reducing NA activity to 5% and 12%, respectively.

**Figure 4:**
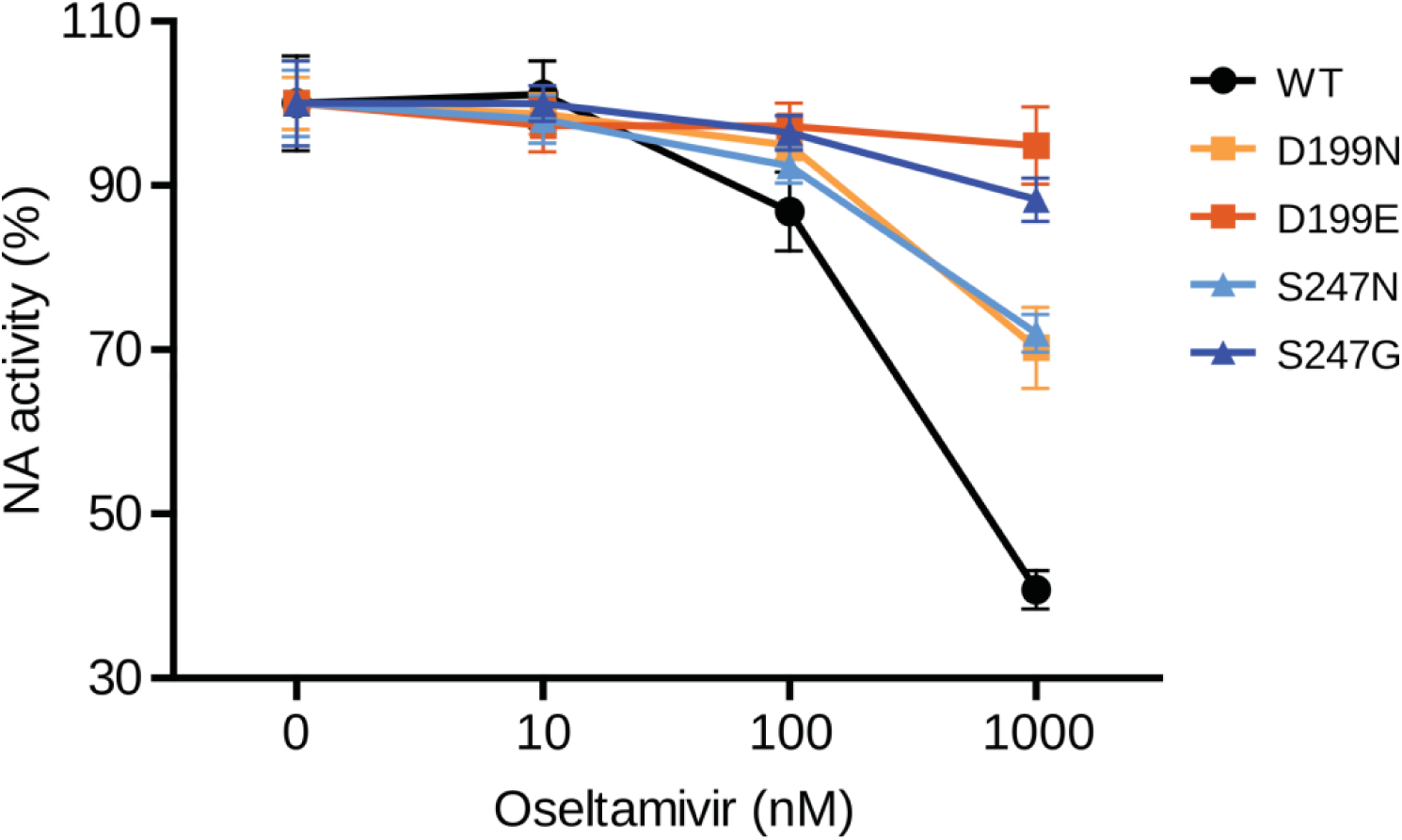
NA activity of pH1N1 viruses in the presence of oseltamivir. NA activity of wt and four NA mutant pH1N1 viruses (D199N, D199E, S247N and S247G) was determined by a fluorescence-based assay using the fluorogenic substrate 4-MU-NANA in the presence of indicated concentrations oseltamivir or mock treated. The mock treated controls of each virus were set to 100 %. Error bars are presented as standard deviation. Shown are the results of three independent experiments performed in triplicates.

In pH1N1/NA, patches 2 and 7, which also include the sweep-related site 117 and 248 ^16^, respectively, and patch 9 overlap or are vicinal to the active center of NA^75-77^. For H3N2/NA, conversely, this was the case for patch 2, which includes active site 151. The patches in NA without links to resistances or immune responses could be relevant for host adaptation, i.e. for establishing its catalytic function in the human host, to establish cleavage activity on an altered substrate range, as pH1N1 viruses had many more patches than H3N2 viruses.

### Host adaptation of the RNP proteins

Reassortant influenza viruses with segments of animal influenza viruses that newly establish in the human population, such as pH1N1, likely require further adaptive changes for fine-tuning the efficiency of replicating and spreading within the human host. NP and the polymerase proteins have important roles in host adaptation^49,78^. Notably, NP had entirely different patch sites in pH1N1 than in H3N2. In H3N2, NP is highly conserved with only one patch under positive selection (450, 451, 453), which does not appear in pH1N1. In pH1N1/NP, the RNA binding pocket surrounds patch 5, indicating a role in fine-tuning the binding of host RNA (Figure 5D)^79^. We also found two patches close to known mammalian adaptation sites: site 100 in patch 2^16,80,81^ and site 319 next to patch 6^82^. Site 319 is also in proximity to sites 373 and 100 implicated in selective sweeps (Figure 5D). The region including patch 2 and patch 6 thus may well be implicated in human adaption in influenza A viruses (Figure 2E), especially, because they have the largest average *dN/dS* value (~1.8) among all NP patches (Supplementary Table 6). Further, of three NP sites (100, 283, 313) relevant for escape recognition by intracellular restriction factor MxA, 313 is located in a beta-sheet adjacent to patch 6 and in vicinity of patch 1^83,84^. Patches 3 and 4 are located in a region of NP with no known relevance for host adaptation.

**Figure 5:**
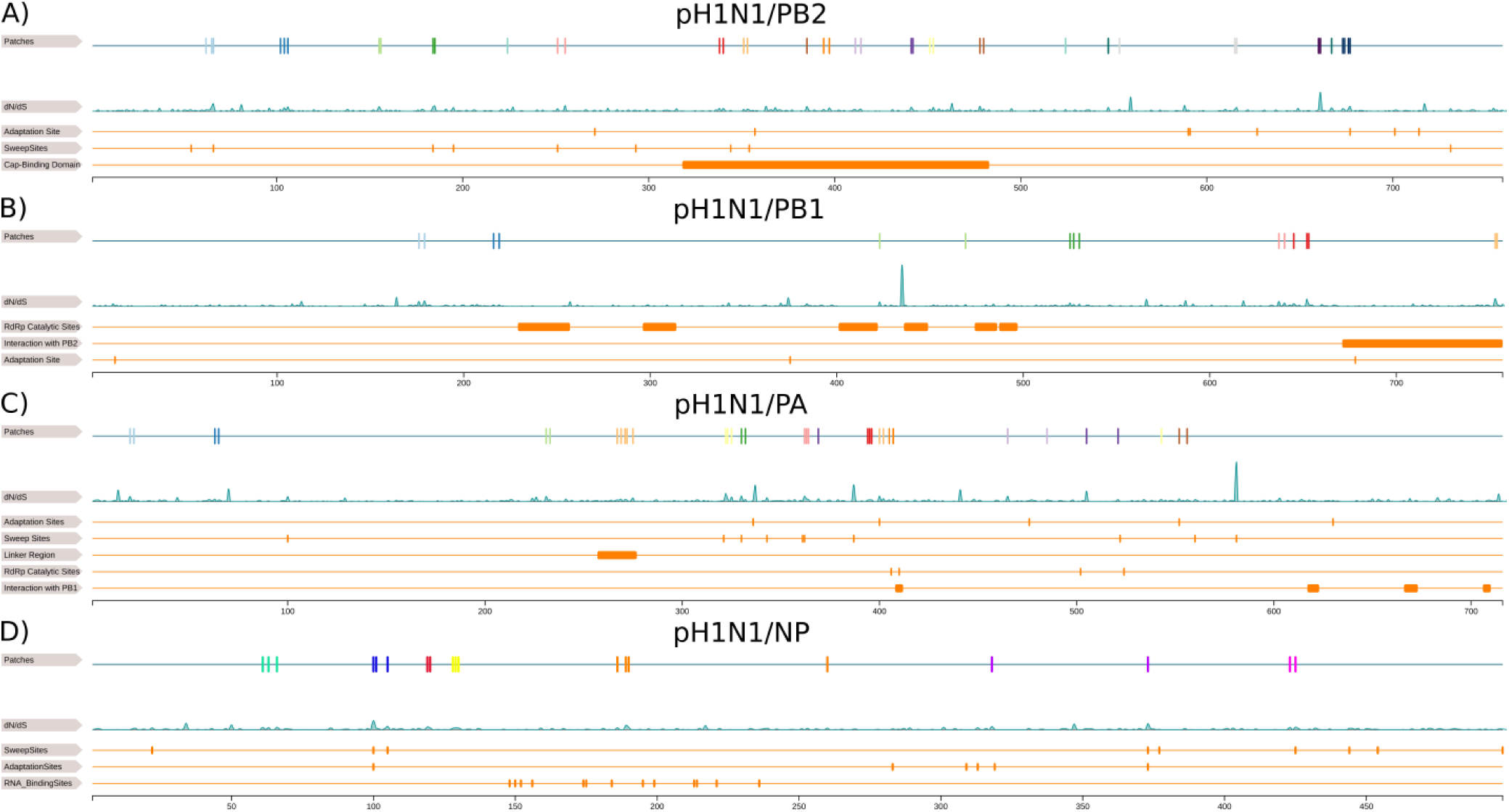
Linear representation of pH1N1 RNP proteins. This figure illustrates the overlap of patch sites with known functional sites in pH1N1 RNP proteins PB2 **(A)** PB1 **(B)** PA **(C)** and NP **(D)**. In each subfigure, the first row is a linear representation of patches and the second row shows the *dN/dS* distribution of the protein. For PB2 **(A)**, we compare with adaptation sites^49^, sweep-related sites^16^, cap binding domain^49^. For PB1 **(B)**, we compare with RdRp catalytic sites^49^, interactions sites with PB2^49,88^ and adaptation sites^89,90^. For PA **(C)**, we compare with adaptation sites ^49^, sweep-related sites ^16^, the linker region of PA^49,87^, RdRp catalytic sites^49^ and interactions sites with PB1^49,87^. For NP **(D)**, we compare with sweep-related sites^16^, adaptation sites^16,80–82^ and RNA binding sites^79^.

Alterations in the polymerase complex proteins PB2, PB1 and PA can determine the host range and confer the ability to infect and replicate in humans. In PB2, site 627 changes when adapting to mammalian hosts and an E to K substitution at this site enhances polymerase activity in influenza A viruses in mammals, while site 701 enhances the pathogenicity and transmissibility of pH1N1 viruses^49^. In pH1N1 viruses, patch 17 includes the adaptation site 677 and patch 6 lies next to adaptation site 357 (Figure 5A). Different from H3N2, for which we could not link patches to adaptation sites, likely because H3N2 is already adapted to the human host, pH1N1 viruses show signs for further host adaptation in PB2. When comparing patch sites with the nine sweep-related sites for pH1N1, three are located within a patch (66 in patch 1, 184 in patch 4 and 251 in patch 5) and one is localized in proximity to a patch (354 to patch 7) (Figure 5A). Remarkably, there are seven patches (6 to 12) in pH1N1 viruses and two patches (3 and 4) in H3N2 viruses within the cap-binding domain (318–483)^49^, indicating that viral adaptation is related to the establishment and the enhancement of transcription activity of PB2.

The PA subunit includes known host adaptation sites 336, 400, 476, 552 and 630, of which we could link site 400 to patch 7, 552 to patch 12 and 336 to patch 4, 5, 10 and 11 in pH1N1 and 552 to patch 6 in H3N2^49^. Interestingly, in pH1N1/PA, site 336 is the central residue of an adaptive cluster including sweep-related sites 321 (proximal to patch 11, average *dN/dS* value (~1.7)), 330 (patch 4, average *dN/dS* value (~1.4)), 361 (patch 5, average *dN/dS* value (~1.2)) and 362 (proximal to patch 5) (Figure 2F&3C)^16,85^. In addition, patch 10, which excludes a sweep-related site, is vicinal to site 336 and has an average *dN/dS* value of ~1.9 (Figure 2F). Other than for H3N2 viruses, for which we detected a single patch next to 336 (patch 2), an intensified signal of positive selection in this region in pH1N1 viruses emphasizes its relevance for further viral adaptation to the human population. For pH1N1, patch 11 suggest further adaptation of polymerase activity, as it is located next to sweep-related site 321, which is known to increase viral polymerase activity^16,86^. The linker region of PA (257-277) is a crucial in the PA and PB1 interaction^49,87^ and includes patch 7 in pH1N1 viruses, but none in H3N2 viruses. This patch has a very strong signal with a 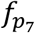 value of 1.0 (Supplementary Table 6). As these segments have different evolutionary origins in the reassortant pH1N1 viruses (PB1 originating from a human H3N2; PA of avian origin, both intermediate in swine), this indicates a further fine-tuning of their interactions. We furthermore detected three patches (patch 7, patch 9 and patch 10) in pH1N1 viruses and one (patch 4) in H3N2 viruses in proximity to the RNA-dependent RNA polymerase activity sites (406, 410, 502 and 524)^49^.

The polymerase subunit PB1 has the fewest number of patches of the polymerase subunits. Patch 4 in H3N2 viruses and patch 7 in pH1N1 viruses overlap with the C-terminal region (671–757), which maintains tight inter-subunit contact with PB2 (1-35)^49,88^. Also for pH1N1, both segments have different evolutionary origins in the reassortant viruses, with PB2 originating more recently from an avian lineage circulating intermittently in swine, indicating a fine-tuning of their interactions^49^. Of three known host adaptation sites (13, 375 and 678), merely patch 2 in H3N2 includes site 375, while no patch of pH1N1 included these sites, which include typical mammalian adaptation residues^89,90^. As pH1N1/PB1 previously circulated in the human population for 30 years and re-entered the human population after circulating for 10 years in swine, the segment is likely already adapted to mammals and humans^5^. The centrally located RNA-dependent RNA polymerase activity motifs of PB1 are highly conserved^49^, with patch 3 in H3N2 viruses and in pH1N1 viruses located in proximity (Figure 5B).

## Discussion

It is well established that the surface antigens of human influenza A viruses evolve under positive selection to escape immune recognition, in an ongoing co-evolutionary arms race with the human adaptive immune response. However, much less is known about the relevance of other proteins in this co-evolutionary arms race and which other factors shape the evolutionary trajectories of these viruses. There is emerging evidence that the pH1N1 virus since 2009 also has acquired changes improving its adaptation to the human host^15,81,86^. The recent introduction of the pH1N1 virus into the human population allows to study the evolutionary forces acting on these viruses in comparison to the H3N2 viruses, which have circulated for almost five decades. To analyze these effects, we searched for protein regions under positive selection from all available historic sequences and protein structures for both influenza A subtypes with a refined computational method, including both structural proximity and the strength of site-wise positive selection into the inference process. While there are studies focusing on selective measurements per protein sites^33-35^, to our knowledge, this study is the first to describe protein regions relevant for adaptation of influenza viruses for almost all viral proteins and a comparative analysis of their functional importance.

Contrary to expectations, the HA1 subunit of HA does not have the largest number of patches or sites under positive selection and was ranked behind NA, M2, NS1 and NS2^34^, an effect becoming more pronounced when analyzing the entire sequence data across all seasons, rather than smaller sequence collections per season. Most BCE sites overlap with patch sites in H3N2 viruses (~70%), while the patches in pH1N1/HA1 marginally overlap with BCE sites. This indicates, as expected, that antigenic evolution over the past decade was not pronounced in pH1N1 viruses, in agreement with a single vaccine strain update for pH1N1 viruses from A/California/7/2009 to A/Michigan/45/2015 being recommended by the WHO for the use in the 2017 southern hemisphere influenza season^91^.

Notably, NS1 is covered most densely with patches of all proteins for both subtypes, suggesting that viral attenuation of innate immune responses is an important factor in the evolution of human influenza viruses, even decades after establishment of the H3N2 virus in the human host, indicating that viral attenuation of host inflammatory immune responses is a continuous process. This is in line with classic evolutionary theory, postulating that in co-evolutionary arms races, both players evolve towards a mitigated state over time causing less severe infections^92,93^. However, recent results have challenged this paradigm, showing the alarming rate of causalities of rabbits infected with myxoma viruses^94^, raising the possibility that the current evolutionary trajectory of down tuning host innate immune defenses for human influenza viruses may also eventually result in an escalation of viral virulence incontrollable by host immune defenses^94^. For pH1N1, patches 3, 7, 9 and 10 include known changes linking to a substantial attenuation of innate immune responses, by restoring NS1-mediated general gene expression inhibition and resulting in less severe inflammatory response after influenza infections^67^. Patches 2 and 7 in H3N2 and patches 6 and 9 in pH1N1 link to CPSF30-binding that initiates inhibition of antiviral IFN-stimulation^65,66^. Further, IFN regulation is altered by amino acid changes at site 196 and 171 in pH1N1^62,69^, as well as 196 and 189 in H3N 2^69,70^, which overlap with patch 6 and 7 in H3N2, as well as patch 9 and 10 in pH1N1. Other sites in patch 9 in pH1N1/NS1 and patch 7 in H3N2/NS1 link to binding host PKR, a critical component of IFN-stimulated defenses^61^. In conclusion, particularly patch 9 in pH1N1/NS1 and patch 7 in H3N2/NS1 may be important for antagonizing IFN responses via CPSF30-binding and other mechanisms in both subtypes.

Overall, we found significantly higher mean *dN/dS* values in most pH1N1 proteins in comparison to H3N2. This was despite the longer time period for which sequence data was available for H3N2 viruses (1968-2016) than for pH1N1 viruses (2009-2016), which we observed to correlate with elevated *dN*/*dS* values across different proteins, suggesting that pH1N1 viruses are not as well adapted to the human population yet as H3N2 viruses are. Accordingly, the number of patches found in PB2, PB1, PA and NP found for pH1N1 viruses exceeds those for H3N2 viruses, and their locations correspond to evolutionary stable areas in H3N2 proteins. For instance, while in H3N2 viruses, NP is largely conserved with only a single patch, in pH1N1 seven patches were found. Patch 2 and 6 are particularly interesting for host adaptation, as they are near known mammalian adaptation sites 100 and 319, respectively^16,80-82^. For PA, the neighboring region around mammalian adaptation site 336 that confers selective advantage to promote polymerase activity in human^85^ overlaps with patches 4, 5, 10 and 11 in pH1N1 viruses, indicating their relevance for human host adaptation. Also for PB2 of pH1N1, patches linking to mammalian adaptation sites (patch 17 and 6) and located in proximity to the cap-binding domain were found, different from H3N2, indicating that recent adaptation of pH1N1 is related to enhanced transcription activity^49^. PB1 of pH1N1 is the protein descending from a recent human lineage, and fewer patches than in the other polymerase subunits were found. Specifically, one patch in the C-terminal region was identified, likely to adjust the interaction with the novel version of PB2 in the reassortant lineage^49^, as PB2 descends from an avian lineage circulating in swine before.

For M2 in both subtypes, we found patches (1 and 2) that include known resistance sites against amantadine and rimantadine, respectively, and multiple currently undescribed sites likely linking to resistances. Similarly, oseltamivir and zanamivir resistances in NA could be linked to patch 2 and 8 in H3N2 and patch 4 and 5 in pH1N1, respectively. In NA activity assays, we could show the relevance of substitutions in patch site 199 and patch site 247 in pH1N1 in conferring resistance to oseltamivir.

The collection of patches and included sites provide an atlas of genetic elements linked to multiple factors predominantly influencing the evolutionary trajectories of human influenza A viruses. We demonstrate the value by confirming the resistance-conferring phenotype of changes at two such sites. The collection provides a rich resource of candidate sites and markers for viral host adaptation, immune evasion, resistance generation and attenuation of host immune responses, which could lead to an improved understanding of the underlying biological processes and more effective monitoring of phenotypes of circulating viral strains.

## Software & Data Availability

The PatchDetection software, figures and tables from this manuscript, and all related data used in this publication are fully available under the Apache License 2.0 at https://github.com/hzi-bifo/PatchDetection.

## Acknowledgements

This work was supported by the Helmholtz foundation. The Heinrich Pette Institute, Leibniz Institute for Experimental Virology is supported by the Free and Hanseatic City of Hamburg and the Federal Ministry of Health. We are grateful to Prof. Thomas Krey for providing the homology models of the pH1N1 and H3N2 polymerase proteins. We thank Hanna Markowsky for excellent technical assistance in performing the NA activity assays.

## Author Contributions Statement

A.C.M conceived the study. A.C.M and T.R.K planned and coordinated the study. A.C.M., T.R.K. and J.L. designed the methodology. T.R.K. and J.L. maintained the data, implemented the software and created results. G.G and S.S.B performed the wet lab experiments. A.C.M., J.L., T.R.K., G.G. and S.S.B analyzed the results. T.R.K., J.L. and S.S.B generated all visualizations. T.R.K., J.L. and A.C.M. wrote the original draft of this manuscript and all authors reviewed the manuscript.

## Competing interests

The authors declare no competing interests.

---

**Figure S1:**
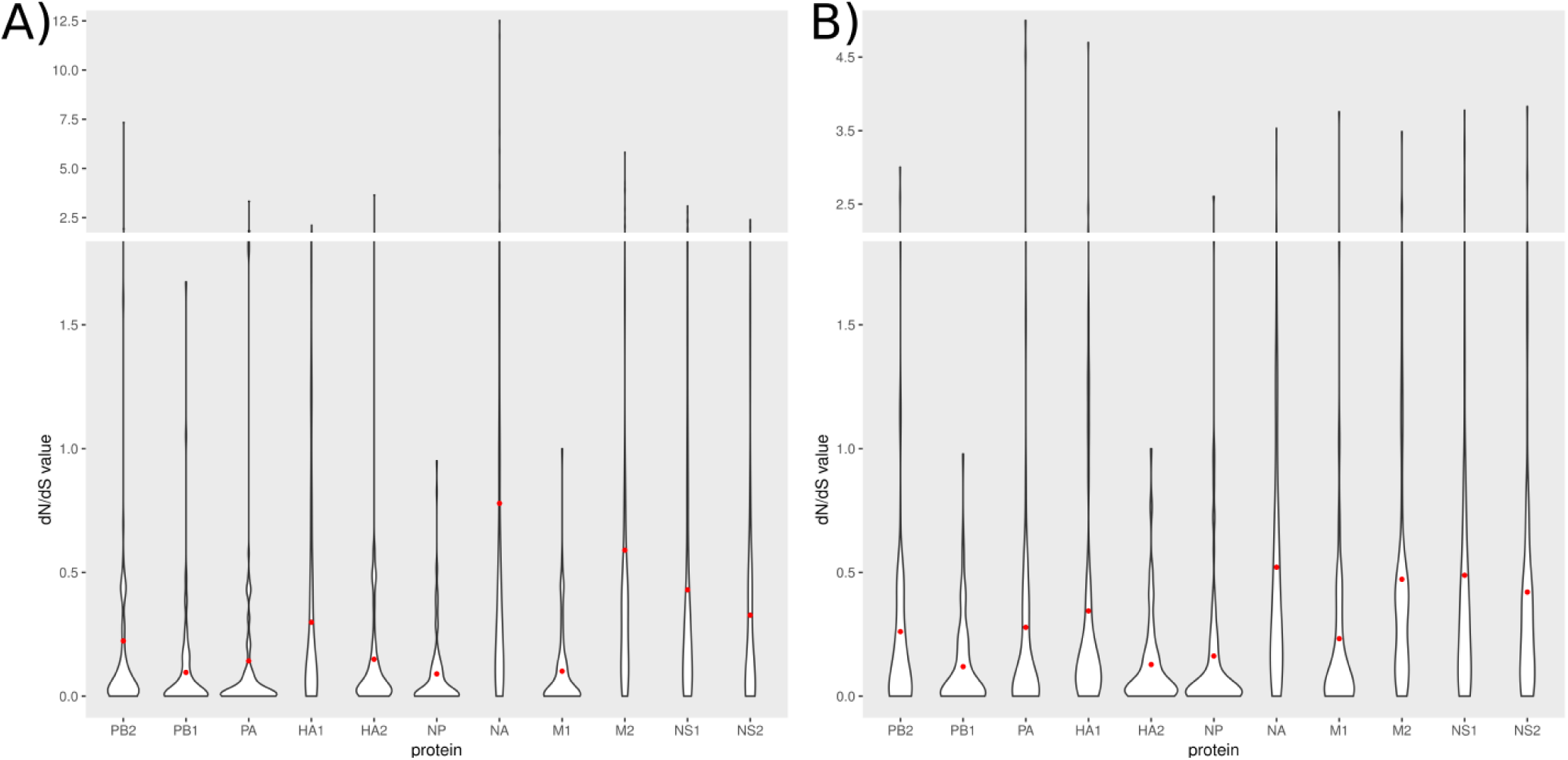
Violin plots of *dN/dS* distribution. This figure illustrates the *dN/dS* distribution in violin plots for H3N2 (A) and pH1N1 (B) viruses. Red dots indicate the mean *dN/dS* value.

**Figure S2:**
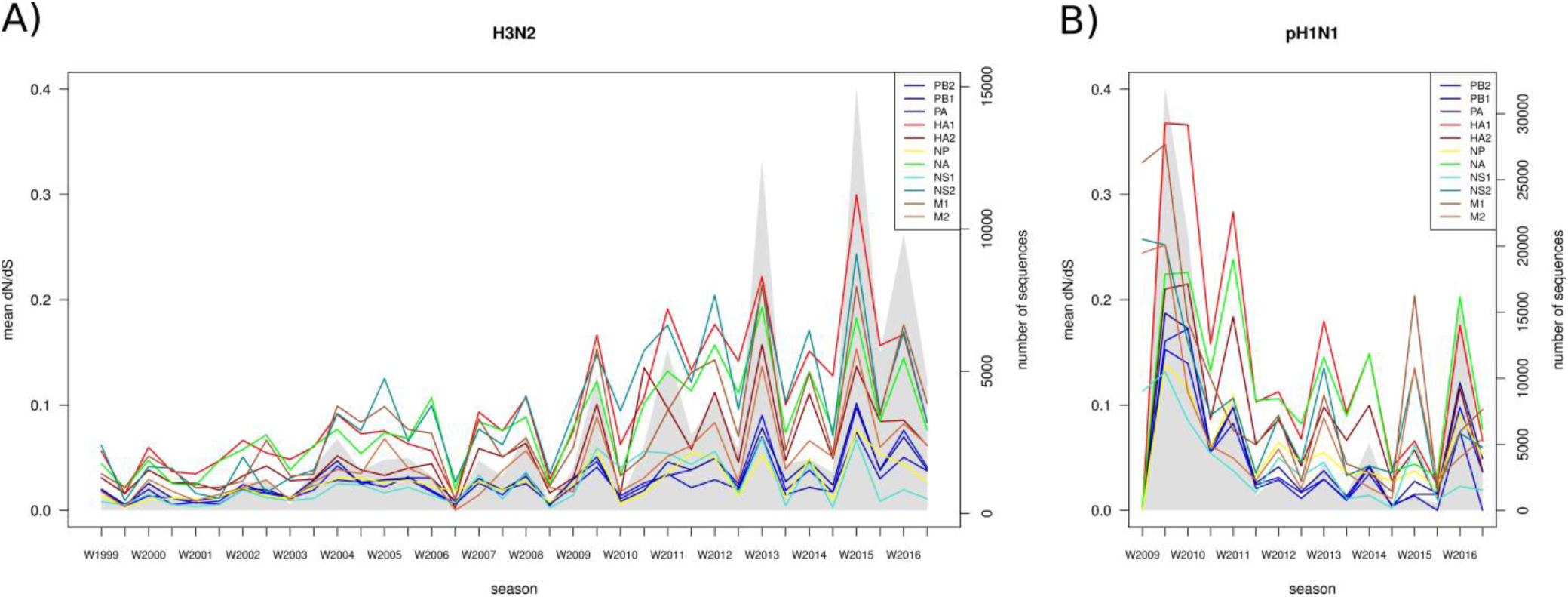
*dN/dS* values per season. We show the average *dN/dS* value per season (left y-axis) for each protein over time for H3N2 **(A)** and pH1N1 **(B)** viruses. The grey area in the background depicts the total number of sequence per season (right x-axis).

**Figure S3:**
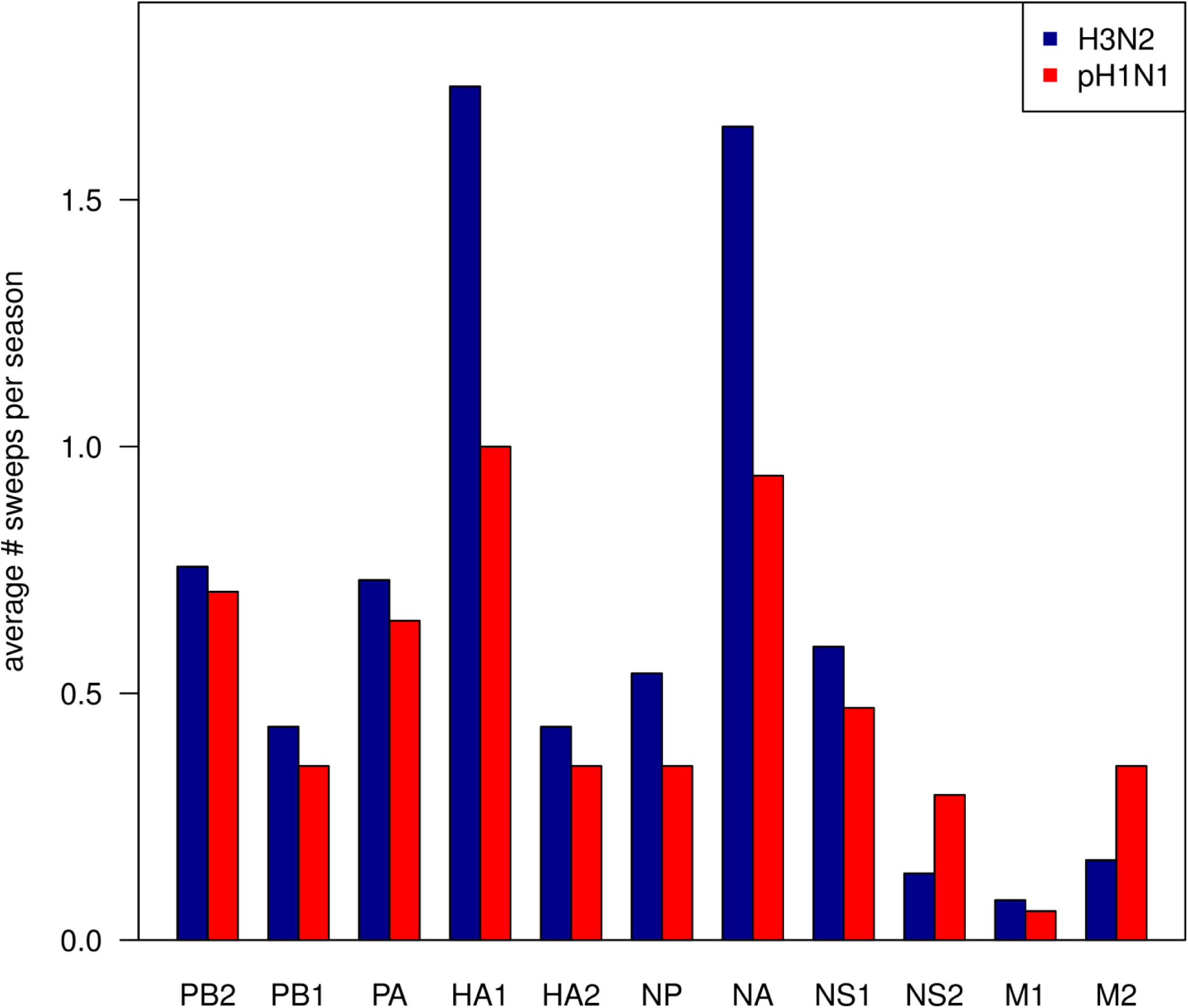
Average number of sweep-related changes. The barplot show the average number of sweep-related changes per season for all proteins of H3N2 **(blue)** and pH1N1 **(red)** viruses.

**Table S1:** Overview of protein modeling for pH1N1 protein. This table provides detailed information about the homology modeling of all pH1N1 proteins.

| Protein | Target Sequence | Template Structures | Template Region | Template Sequence | Sequence Identity | Region | Coverage |
| --- | --- | --- | --- | --- | --- | --- | --- |
| PB2 | A/California/07/2009(H1N1) | 4wsbC | 1-482, 491-741 | A/little yellow-shouldered bat/Guatemala/060/2010 (H17N10) | 67% | 1-759 | 100% |
| PB1 |  | 4wsbB | 1-194, 199-490, 492-647, 650-733, 736-754 |  | 79% | 1-757 | 100% |
| PA |  | 4wsbA | 1-185, 204-714 |  | 72% | 1-716 | 100% |
| HA1 |  | 3m6sA | 1-322 | A/Darwin/2001/2009 (H1N1) | 99% | 1-326 | 99% |
|  |  | 3lzgA | 11-325 | A/California/04/2009(H1N1) | 99% |  |  |
| HA2 |  | 3m6B | 1-170 | A/Darwin/2001/2009 (H1N1) | 98% | 1-175 | 100% |
|  |  | 3lzgB | 1-175 | A/California/04/2009(H1N1) | 98% |  |  |
| NP |  | 3ro5A | 22-71, 91-198, 212-429, 439-479, 485-497 | A/Wilson-Smith/1933 (H1N1) | 92% | 21-498 | 96% |
|  |  | 3ro5B | 22-83, 88-199, 214-246, 248-393404-428, 441-479, 486-491 |  | 92% |  |  |
| NA |  | 4b7QA | 83-350, 352-469 | A/California/07/2009(H1N1) | 100% | 83-469 | 83% |
| M1 |  | 3md2A | 4-67, 77-85, 89-157 | A/California/04/2009 (H1N1) | 100% | 1-157 | 62% |
|  |  | 5cqeA | 1-71, 73-158 | A/Puerto Rico /8/34(H1N1) | 95% |  |  |
| M2 |  | 2n70A | 18, 20-30, 32-49, 51-60 | A/Udorn/307/1972(H3N2) | 81% | 18-62 | 47% |
|  |  | 2kihA | 23-30, 32-49, 51-60 |  | 84% |  |  |
|  |  | 2l0jA | 22-49, 51-62 |  | 83% |  |  |
| NS1 |  | 3m5rB | 79-203 | A/California/07/2009(H1N1) | 100% | 1-219 | 100% |
|  |  | 4opaB | 2-37, 41, 42-49, 85-203 | A/blue-winged teal/MN/993/1980(H6N6) | 84% |  |  |
|  |  | 3m8aA | 1-72 | A/California/07/2009(H1N1) | 100% |  |  |
| NS2 |  | 1pd3 | 64-85, 87-103, 105-116 | A/Puerto Rico /8/1934(H1N1) | 96% | 64-114 | 42% |

**Table S2:** Overview of protein modeling for H3N2 protein. This table provides detailed information about the homology modeling of all H3N2 proteins.

| Protein | Target Sequence | Template Structures | Template Region | Template Sequence | Sequence Identity | Region | Coverage |
| --- | --- | --- | --- | --- | --- | --- | --- |
| PB2 | A/Aichi/2/1968 (H3N2) | 4wsbC | 1-482, 491-741 | A/little yellow-shouldered bat/Guatemala/060/2010 (H17N10) | 67% | 1-759 | 100% |
| PB1 |  | 4wsbB | 1-194, 199-490, 492-647, 650-733, 736-754 |  | 79% | 1-757 | 100% |
| PA |  | 4wsbA | 1-185, 204-714 |  | 70% | 1-716 | 100% |
| HA1 |  | 3hmgA | 1-225, 227-326 | A/Aichi/2/1968 (H3N2) | 100% | 1-328 | 100% |
| HA2 |  | 3hmgB | 1-175 |  | 100% | 1-175 | 79% |
| NP |  | 4x9aA | 21-72, 92-202, 213-396, 403-428, 438-489 | A/Wilson-Smith/1933 (H1N1) | 91% | 21-498 | 96% |
|  |  | 3zdpB | 21-391, 408-498 |  | 92% |  |  |
| NA |  | 3tiaA | 82-469 | A/RI/5+/1957 (H2N2) | 88% | 81-469 | 83% |
|  |  | 2aepA | 82-469 | A/Memphis/31/98(H3N2) | 98% |  |  |
| M1 |  | 1ea3 | 2-158 | A/Puerto Rico/8/1934 (H1N1) | 98% | 1-160 | 63% |
|  |  | 5cqeB | 1-159 |  | 98% |  |  |
| M2 |  | 2ly0A | 22-49, 51-62 | A/Chiba/5/71 (H3N2) | 97% | 18-62 | 47% |
|  |  | 2n70A | 18-60 | A/Udorn/307/1972(H3N2) | 87% |  |  |
| NS1 |  | 1ailA | 1-70 | A/Udorn/307/1972 (H3N2) | 90% | 1-230 | 97% |
|  |  | 4ophA | 2-37, 29-40, 42-203 | A/blue-winged teal/MN/993/1980 (H6N6) | 85% |  |  |
|  |  | 3d6rB | 83-202 | A/Duck/Alberta/60/1976 (H12N5) | 73% |  |  |
|  |  | 3ee9A | 84-205 | A/Udorn/307/1972(H3N2) | 94% |  |  |
| NS2 |  | 1pd3 | 64-85, 87-103, 105-116 | A/Puerto Rico /8/1934(H1N1) | 96% | 63-116 | 45% |

**Table S3:** Comparison of mean *dN/dS*. As a sanity check, we compared our results with the mean *dN/dS* value from the Suzuki-Gojobori counting approach implemented in HyPhy SLAC for all proteins.

|  | **H3N2** |  | **pH1N1** |  |
| --- | --- | --- | --- | --- |
|  | **dN/dS** | **HyPhy SCLAC** | **dN/dS** | **HyPhy SCLAC** |
| **PB2** | 0.223 | 0.283 | 0.260 | 0.480 |
| **PB1** | 0.096 | 0.204 | 0.119 | 0.220 |
| **PA** | 0.142 | 0.212 | 0,277 | 0.455 |
| **HA1** | 0.298 | 0.522 | 0,344 | 0.506 |
| **HA2** | 0.148 | 0.206 | 0.127 | 0.231 |
| **NP** | 0,089 | 0.127 | 0,162 | 0.269 |
| **NA** | 0,778 | 0.688 | 0,520 | 0.950 |
| **M1** | 0,100 | 0.208 | 0,232 | 0.350 |
| **M2** | 0.589 | 1.812 | 0,472 | 1.242 |
| **NS1** | 0.429 | 0.589 | 0,488 | 0.977 |
| **NS2** | 0.327 | 0.774 | 0,420 | 0.611 |

**Table S4:** Kolmogorov–Smirnov-test. We applied the Kolmogorov–Smirnov-test (KS-test; *H*_0_ *: dN/dS* distribution of pH1N1 protein is smaller than *dN/dS* distribution of H3N2 protein) to all proteins individually and showed that the global *dN/dS* distributions of pH1N1 proteins were significantly larger than those of H3N2 proteins, except for HA2, NS1 and M2.

| **protein** | **p-value** |
| --- | --- |
| **PB2** | $3 * 10^{-10}$ |
| **PB1** | $2.7 * 10^{-7}$ |
| **PA** | $4.97 * 10^{-9}$ |
| **HA1** | $2.38 * 10^{-2}$ |
| **HA2** | $6.005 * 10^{-1}$ |
| **NP** | $1.46 * 10^{-3}$ |
| **NA** | $1.33 * 10^{-2}$ |
| **M1** | $65 * 10^{-5}$ |
| **M2** | $6.89 * 10^{-1}$ |
| **NS1** | $1.48 * 10^{-1}$ |
| **NS2** | $15.34 * 10^{-4}$ |

**Table S5:** Patch site frequency per protein. We list the ratio of patch sites to protein sites of the protein model for each protein individually.

|  | **sH3N2** | **pH1N1** |
| --- | --- | --- |
| **PB2** | 2.503 % | 5.270 % |
| **PB1** | 1.453 % | 2.113 % |
| **PA** | 2.094 % | 4.748 % |
| **HA1** | 16.158 % | 15.853 % |
| **HA2** | 1.142 % | 0 % |
| **NP** | 0.602 % | 3.815 % |
| **NA** | 7.675 % | 8.955 % |
| **M1** | 0 % | 1.587 % |
| **M2** | 11.340 % | 12.371 % |
| **NS1** | 23.043 % | 27.397 % |
| **NS2** | 0 % | 0 % |

**Table S6:**
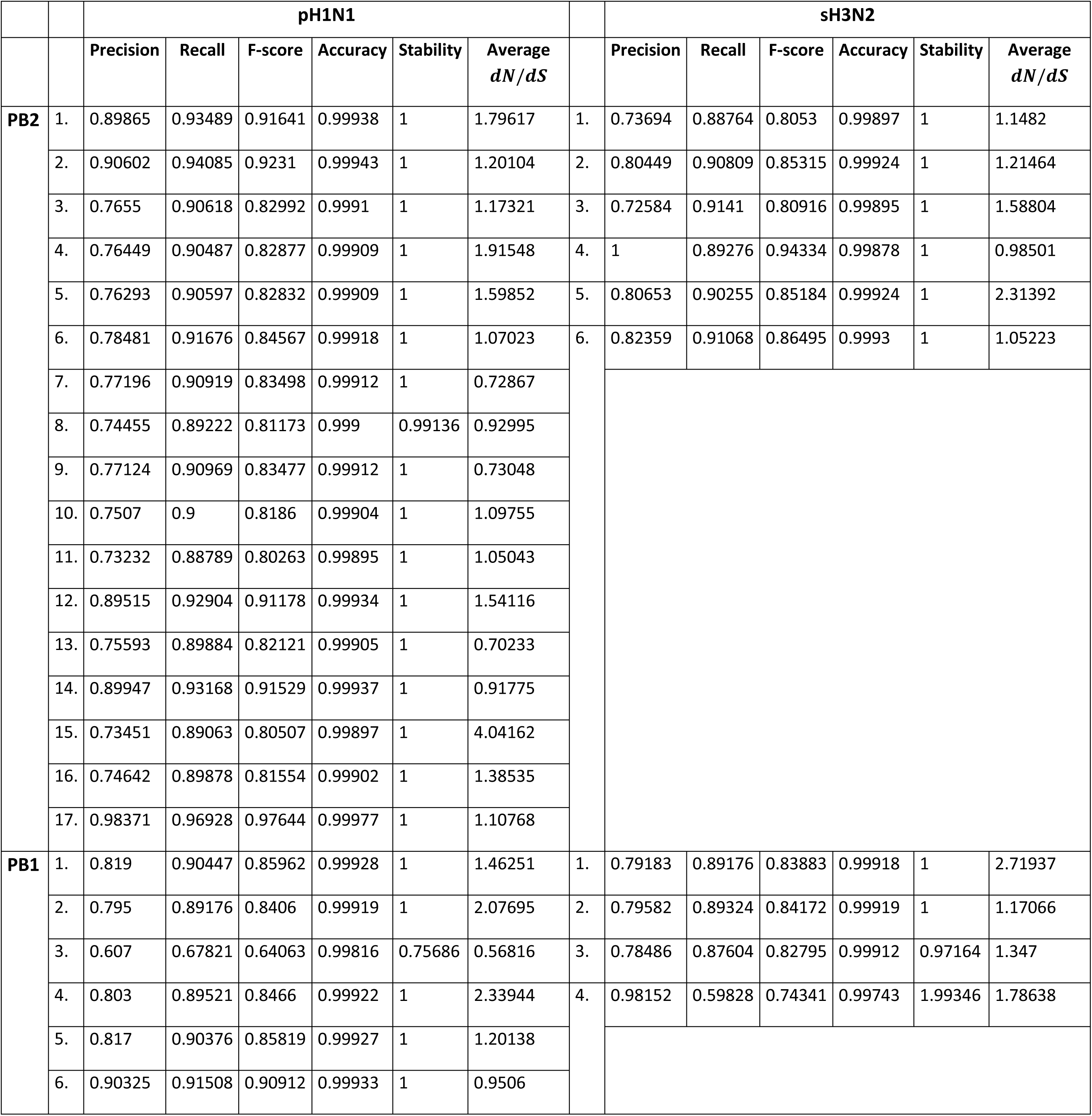
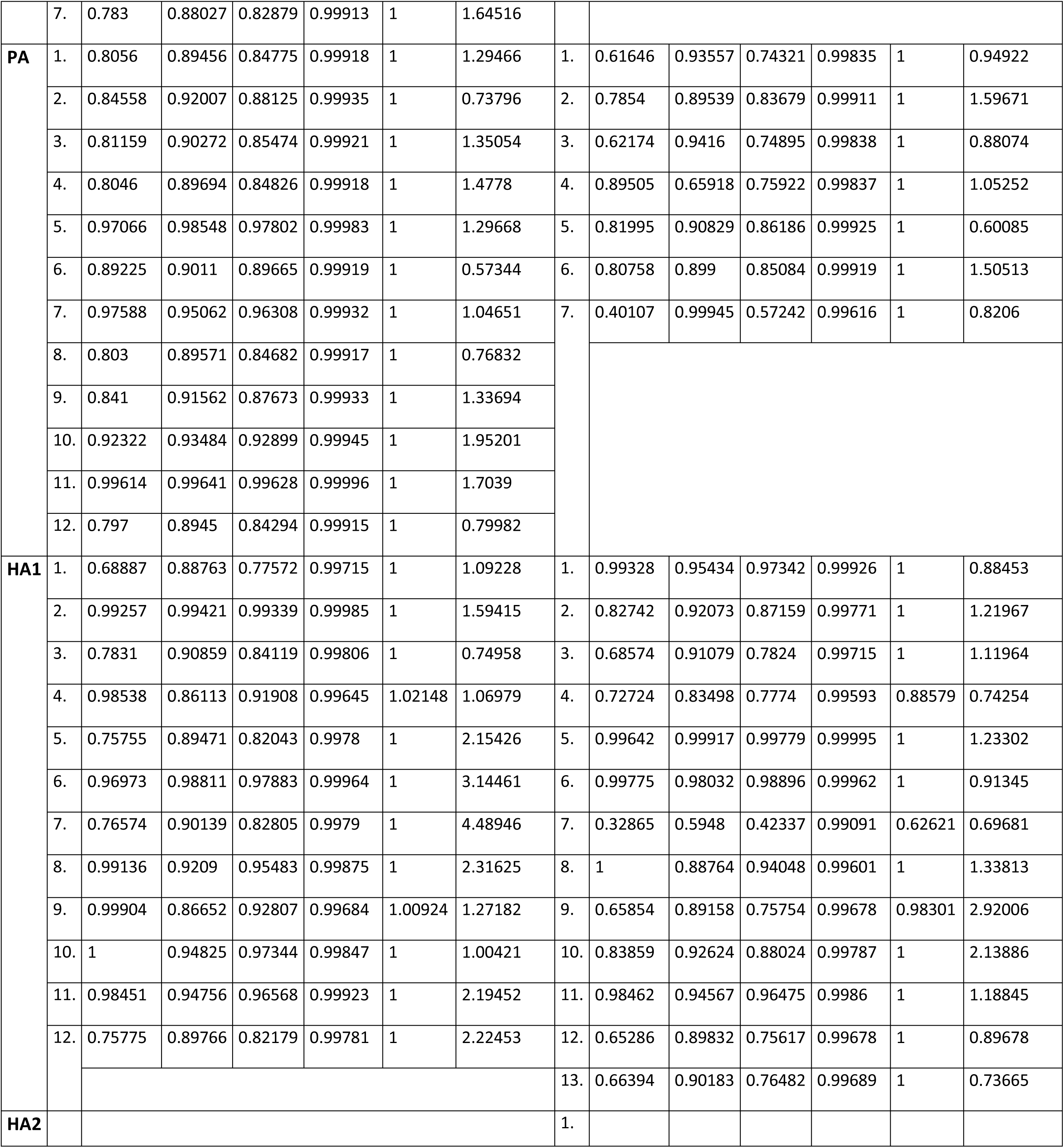
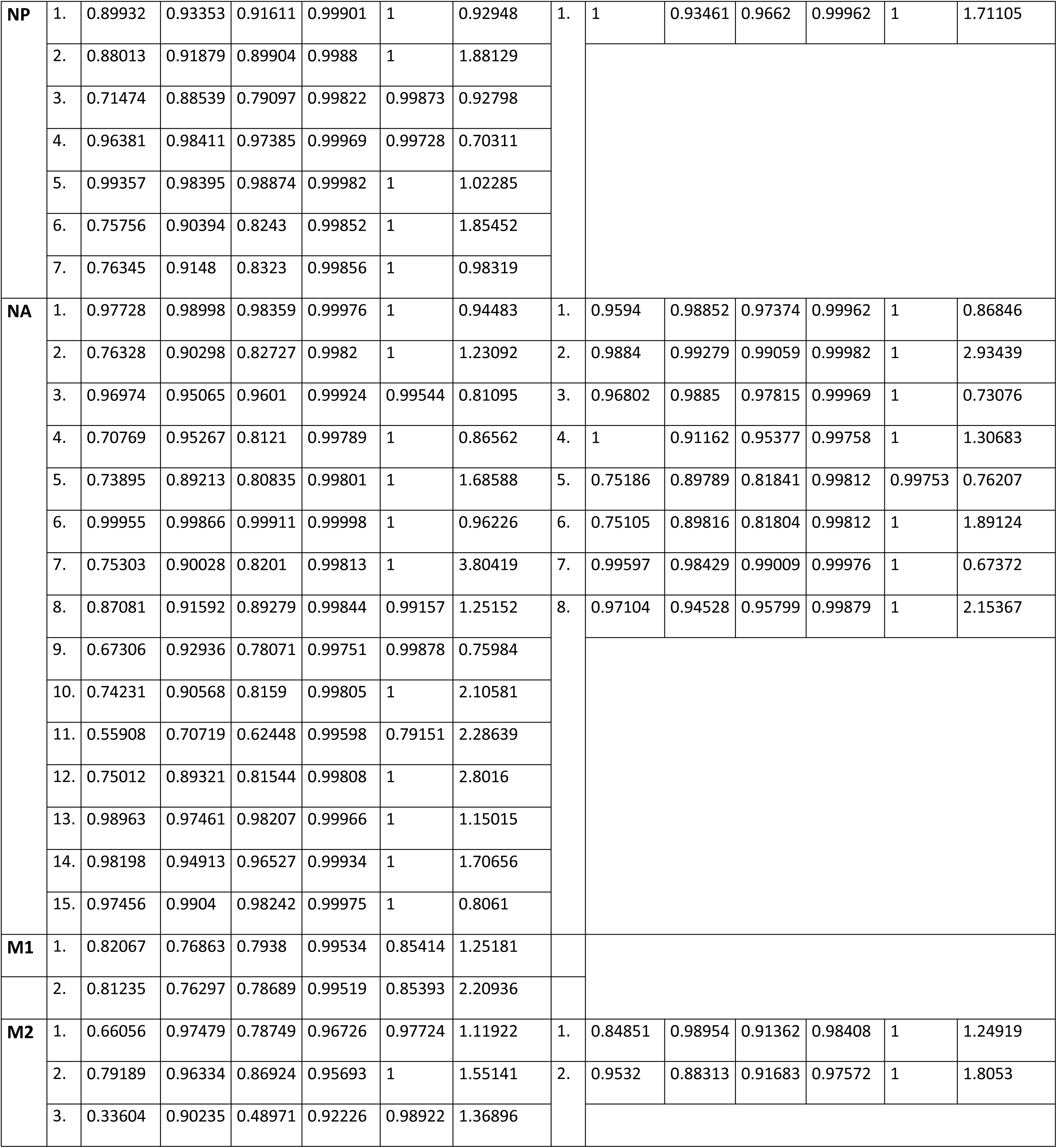
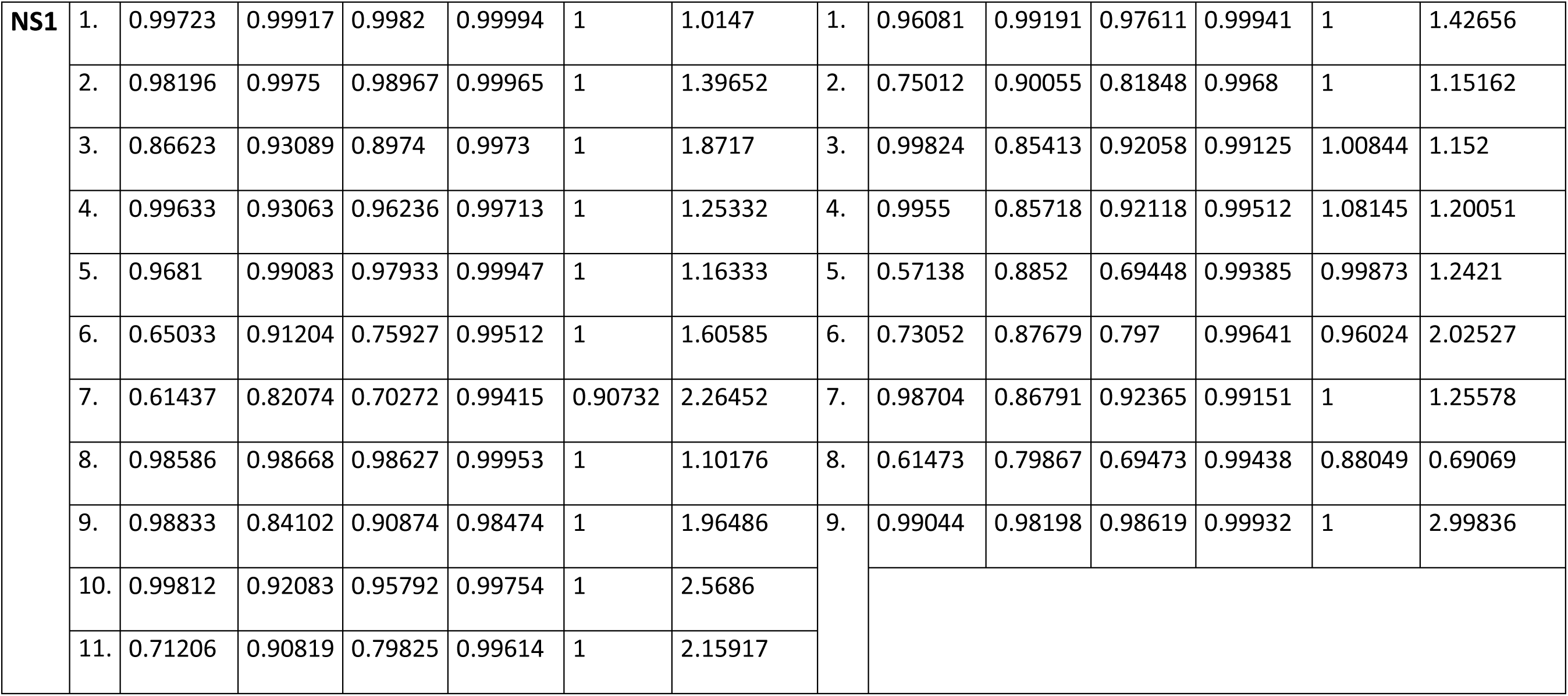
Overview of all quality measurements per patch. This is a detailed overview providing the precision, recall, F-score, stability and the average *dN/dS* of all patches.

